# Host oxidative stress primes mycobacteria for rapid antibiotic resistance evolution

**DOI:** 10.1101/2025.11.19.689367

**Authors:** Evan Pepper-Tunick, Vivek Srinivas, Fred D. Mast, Song Li, Sagan Russ, Weston Hanson, Amy D. Zamora, Wei-Ju Wu, Matthew Silcocks, Dang Thi Minh Ha, Sarah J. Dunstan, Thuong Nguyen Thuy Thuong, Serdar Turkarslan, John D. Aitchison, Mario L. Arrieta-Ortiz, Nitin S. Baliga

## Abstract

The rapid emergence of multidrug-resistant *Mycobacterium tuberculosis* (Mtb) threatens global TB control, yet the mechanisms enabling rapid evolution of drug resistance in Mtb remain poorly understood. Here we reveal that pre-existing mutations in oxidative stress response genes create permissive genomic backgrounds that accelerate high-level isoniazid resistance (INH^R^) without fitness costs, challenging the paradigm that resistance mutations always precede their fitness compensatory adaptations. Using *M. smegmatis* mc^2^155 (Msm) as a model, we show that brief exposure to sublethal INH (2× IC_50_) enriches for "low-level resistance and tolerance" (LLRT) mutants in a single step. These LLRT mutants, particularly those with *ohrR* loss-of-function mutations, acquire high-level resistance (> 500× IC_50_) at 6-fold higher rates than wildtype, primarily through otherwise-deleterious mycothiol biosynthesis mutations that become tolerable in the oxidative stress-buffered background. Crucially, we demonstrate that sublethal oxidative stress alone, mimicking host immune pressure, nearly tripled the rate of INH resistance evolution in Msm. Bayesian analysis of 1,578 clinical Mtb isolates from Vietnam confirmed that mutations in oxidative stress response genes were significantly associated with the emergence of INH^R^ strains (*p-value* = 1.09×10^-7^). Independently, reanalysis of genome-wide CRISPRi screens revealed that the OSR network and high Bayes probability genes are functionally associated with treatment escape and survival with multiple antibiotics, including isoniazid, rifampicin, ethambutol, bedaquiline, vancomycin, clarithromycin, linezolid, and streptomycin. Our findings that host-imposed oxidative stress and inadequate drug penetration may synergistically prime Mtb populations for rapid resistance evolution suggest that targeting pre-resistance mechanisms, such as oxidative stress defenses, could help slow the emergence of antibiotic resistance in tuberculosis.

## Introduction

The rise in drug-resistant *Mycobacterium tuberculosis* (Mtb) infections, the bacterium responsible for tuberculosis (TB), is a growing global health crisis (*Global Tuberculosis Report 2023*), creating an urgency to understand the mechanisms driving rapid gain of antimicrobial resistance in mycobacteria. Because antibiotics target essential functions, such as cell wall synthesis, DNA replication, transcription, translation, or metabolism, resistance conferring mutations often carry a fitness cost typically resulting in a decreased growth rate of the microbe in the absence of the selective pressure (Paulander et al., 2009; Hall & MacLean, 2011; Rajer et al., 2022). Despite the associated fitness cost, the introduction of new antimicrobial compounds is often immediately followed by the emergence of clinically resistant strains (Andersson et al., 2020; Cantón & Morosini, 2011). Compensatory adaptations can ameliorate the fitness cost of the resistance conferring mutations through a number of mechanisms, including epistatic interactions, gene duplications, the use of alternative pathways to reduce the need of the drug target, and many others (Levin et al., 2000; Björkman et al., 2000; Arrieta-Ortiz et al., 2022). In the presence of a strong selective pressure, resistant cells will outcompete their susceptible cousins and eventually dominate the population, especially if selective pressures remain constant. However, in natural environments, including both clinical and agricultural settings, the strength and type of forces that determine which mutant alleles are selected for or against usually fluctuate. Bacterial pathogens may be exposed to low concentrations of antibiotics for extended periods of time, allowing low-cost mutations that give some survival advantage an opportunity to be enriched and subsequently selected (Baquero et al., 1998; Frimodt-Møller et al., 2018; Gullberg et al., 2011; A. Liu et al., 2011; Sandegren, 2014). For example, *in vivo* and *in silico* pharmacokinetic studies have suggested that inadequate antibiotic penetration and accumulation into the granulomas of tuberculosis patients can be common, and may occur for a number of reasons, including granuloma heterogeneity or treatment non-compliance (Cicchese et al., 2020; Cronan, 2022; Day et al., 2023; Lin & Liao, 2015). The survival advantage conferred by mutations that precede full resistance (pre-resistance mutations) may be subtle, generating low-level resistance, tolerance, or other phenotypes that ultimately buy these cells time to propagate and acquire full, clinical resistance conferring mutations (Levin-Reisman et al., 2017). Despite significant progress in identifying and characterizing bacterial antibiotic-resistance mechanisms, we still do not fully understand how sub-inhibitory antibiotic exposure or host-induced stress triggers phenotypic resistance or tolerance. These conditions may also enrich pre-resistance mutations — variants that occur at relatively high frequencies because they carry low fitness costs.

Here, we report findings from an integrated strategy that leverages laboratory evolution using *M. smegmatis* (Msm) as a model mycobacterial species with parallel analysis of genotypic and phenotypic (antibiotic susceptibility) characteristics of clinical Mtb strains to identify pre-resistance mutations that precede and ultimately potentiate rapid gain of high-level resistance to the frontline TB drug isonicotinic acid hydrazide (isoniazid: INH). Our findings show that the mitigation of oxidative stress is a hallmark feature of pre-resistance in *Mycobacterium spp.*, and that mutations that enhance the ability to neutralize reactive oxygen species (ROS) create fertile grounds for selecting mutations conferring high-level INH resistance, without fitness tradeoff. Contrary to prior interpretations that compensatory adaptations appear later to ameliorate the cost of antibiotic resistance, our findings show that these adaptive changes are due to pre-resistance mutations that appear earlier, and accelerate the gain of high-level resistance-conferring mutations without the associated fitness tradeoff. Importantly, we discovered that brief pre-exposure of Msm to sub-inhibitory levels of ROS, to mimic host-induced stress, nearly tripled the rate of gain of high-level resistance to INH, and furthermore, that genes in the Mtb oxidative stress response (OSR) network were functionally associated in CRISPRi screens with escape and survival from treatment with multiple frontline drugs. These findings help explain why *M. tuberculosis* rapidly acquires clinical resistance to new antibiotics and point to strategies that could slow the emergence of drug resistance.

## Results

### Brief one-step exposure to low-dose antibiotic enriches sub-populations of Msm with low-level resistance and tolerance to INH with no fitness trade-off

We investigated the consequence of brief exposure to a low-dose of antibiotic, as experienced by Mtb bacilli within a granuloma due to inadequate penetration of the drug, by subjecting 8 replicate lines of log-phase Msm (mc^2^155) to 2× IC_50_ (8.0 µg/mL) INH for 16 hours. Following the brief treatment, culture aliquots were plated on 7H10 agar with 2× IC_50_ INH and screened with ScanLag (Levin-Reisman et al., 2010) to assess phenotypic heterogeneity of putative low resistance sub-populations of Msm (**Figure S1**). Altogether 255 colonies across the eight replicate lines clustered into three groups (**Figure 1A**) based on PCA and k-means analysis of growth rate (**Figure 1B**), time of appearance (**Figure 1C**), and maximum colony size. Fifty-five colonies (at least six colonies from each replicate line), representative of phenotypic diversity across the three clusters, were characterized further to determine change in fitness in absence of antibiotic and level of resistance to INH (fold change in IC_50_) relative to the wildtype ancestor. While there were no significant differences in average fitness and IC_50_ of isolates from each cluster relative to the wildtype (**Figures 1D-E**), at least 40 isolates exhibited low-level resistance (≥1.1× fold change IC_50_ with respect to wildtype) with little-to-no fitness tradeoff (**Figure 1F**). In summary, these findings demonstrate that low-level INH resistant sub-populations were enriched with brief exposure to low-dose antibiotic, and that these mutants co-exist within a larger naïve wildtype mycobacterial population, even in the absence of the antibiotic.

**Figure 1:**
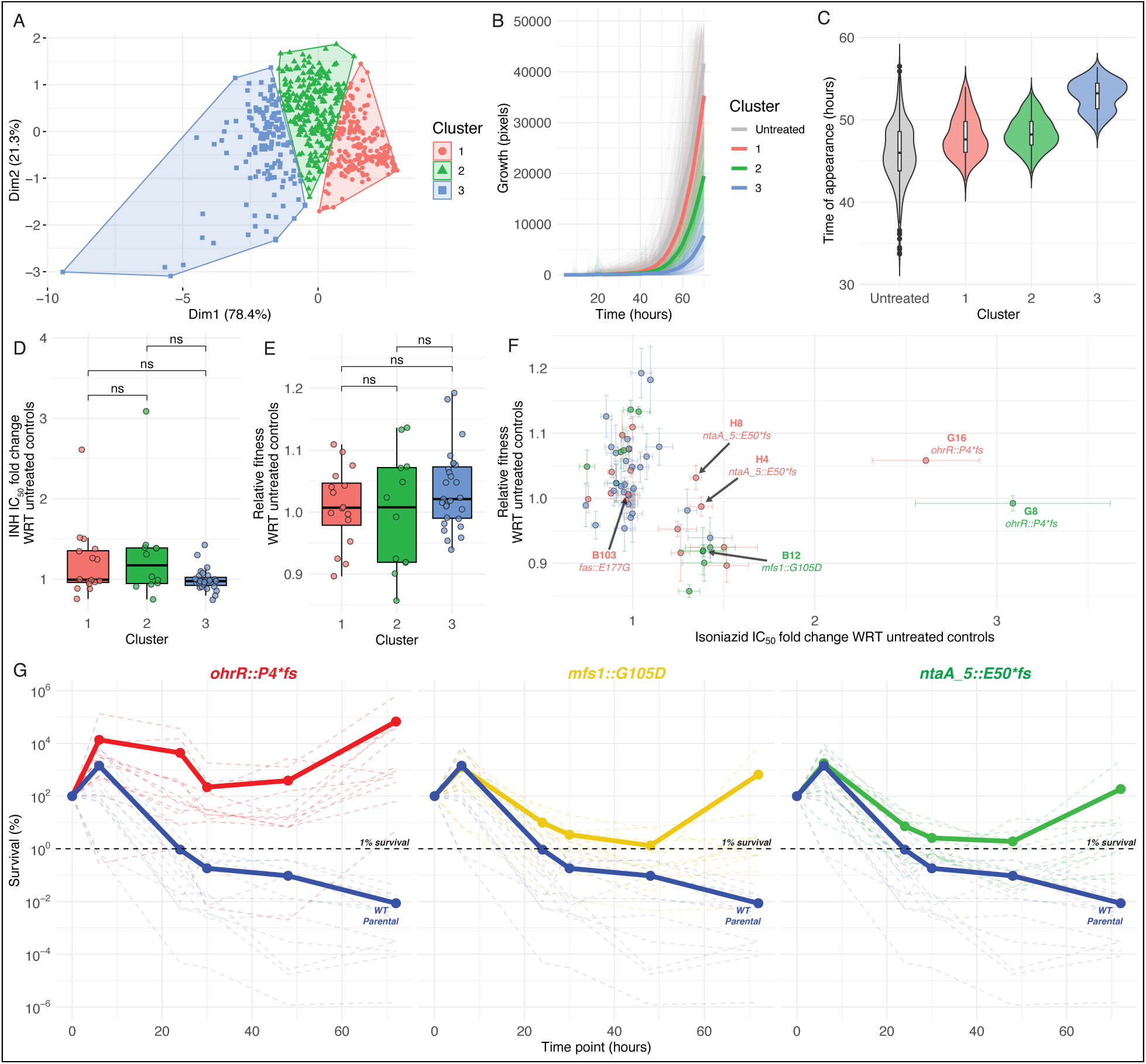
Brief treatment with 2× IC_50_ isoniazid enriches subpopulations with low-level resistance and tolerance, and no fitness tradeoff. **(A)** Principal component analysis of the time of appearance, maximum growth rate, and maximum size of each colony on 7H10 agar with 2× IC_50_ INH following 16 hours-exposure to 2× IC_50_ INH in broth. Growth curves **(B)** and time-of-appearance **(C)** on antibiotic-free agar for colonies from each of the three clusters in **A** and control (“untreated”) Msm cultures. Each solid line represents the mean growth curve profile of colonies within each cluster (untreated and treated + clustered). Fold change in IC_50_ of INH **(D)** and relative fitness **(E)** for isolates in each of the three clusters. **(F)** IC_50_ of INH and fitness in antibiotic-free growth medium for at least 12 isolates from each cluster (total n = 55) relative to untreated controls across four replicates (from duplicate cultures across 2 independent experiments). The gray box centered on coordinates (1, 1) represents standard errors in relative fold changes in INH IC_50_ (width of box) and fitness (height of box) across all untreated control isolates (n = 12). **(G)** Time-kill curves with 100× MIC INH (1 mg/mL) for each of the three mutants and the wildtype strain. Solid lines represent average kill curve for 12 replicates of each strain. Statistical significance was evaluated using the Student’s t-test (\**p* < 0.05; \*\**p* < 0.01; \*\*\**p* < 0.001).

Genome re-sequencing and variant analysis suggested that the low-level resistance phenotypes of at least 15% (6/40) of all sequenced isolates may have emerged independently through point mutations in four genes: *ohrR*, *mfs1*, *ntaA_5* and *fas1* (**Table 1**). Specifically, two isolates from the same line had a single base insertion resulting in frameshift at residue 4 (P4* *fs*) of *ohrR* (organic hydroperoxide reductase regulator). Loss of function mutations in OhrR, a transcriptional repressor of the *ohr* gene which encodes an organic hydroperoxide reductase, has been previously associated with low-level INH resistance in Msm (Meireles et al., 2022; Saikolappan et al., 2014). One isolate with a nonsynonymous mutation in the major facilitator superfamily 1 gene, *mfs1* (G105D), exhibited ∼1.8× fold increase in INH IC_50_. MFS transporters, including the Mtb ortholog Rv2994, have been implicated in antibiotic efflux in many bacterial species (Buchmeier et al., 2003; Titgemeyer et al., 2007). Two isolates harbored a frameshift mutation at residue 51 (E50* *fs*) in *ntaA_5*, which encodes a putative xenobiotic compound monooxygenase from a class of enzymes implicated in detoxification of antibiotics, including tetracycline, rifampicin and imipenem (Andersen et al., 1997; L.-K. Liu et al., 2016; Minerdi et al., 2015, 2023; Moore et al., 2005; Reis et al., 2021; Rudra et al., 2018; Volkers et al., 2011; W. Yang et al., 2004). Lastly, a sixth isolate with a nonsynonymous mutation (E177G) in *fas1*, which encodes fatty acid synthase 1 (FAS1), an enzyme required for *de novo* biosynthesis and elongation of fatty acids that ultimately become mycolic acids in the mycobacterial cell wall (Elad et al., 2018). While FAS1 is yet to be associated with antibiotic resistance, the reaction products of FAS1 are reactants for the FAS II-system, mutations in which, including in InhA, are known to confer high-level resistance to INH in Msm and Mtb (Banerjee et al., 1994; Larsen et al., 2002; Machado et al., 2013; Morlock et al., 2003; Rouse et al., 1995).

**Table 1.**
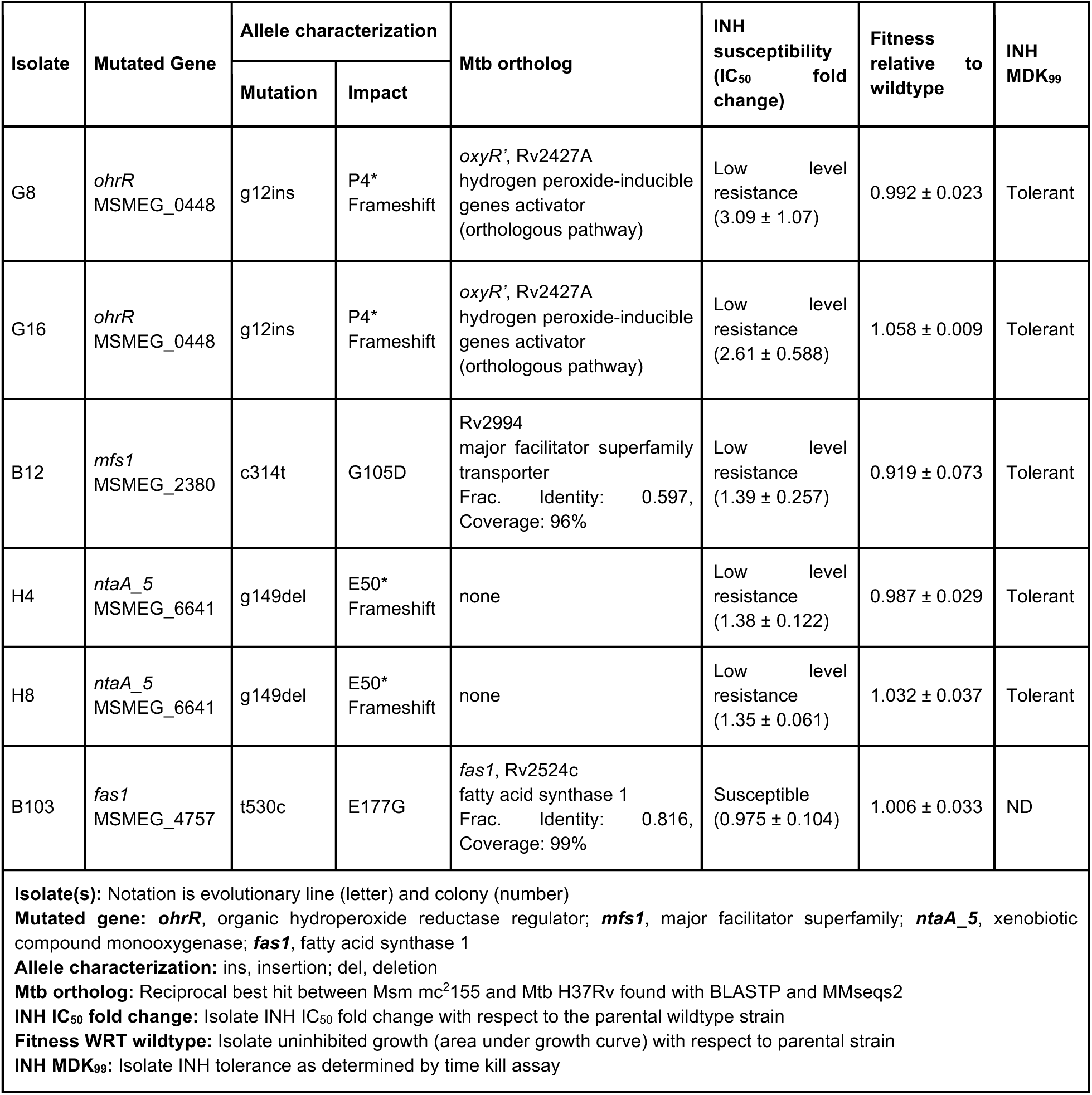
Mutations in three genes confer low-level resistance and tolerance to INH.

Prior work in *E. coli* has demonstrated that cyclic exposures to high-dose antibiotic, with an intermediate counterselection of fitness-compromising mutations, enriches sub-populations with phenotypic tolerance, which is eventually fixed through selection of tolerance-conferring mutations (Levin-Reisman et al., 2017). We performed time kill assays to investigate whether mutants selected with a single short-term exposure to low-dose INH also conferred tolerance to the antibiotic. We added 100× the wildtype minimum inhibitory concentration (MIC) of INH (1 mg/mL) to mid-log phase cultures of the parental wildtype mc^2^155 and each of the three mutants (excluding the *fas1::*E177G mutant), and assayed loss of survival over a 72-hour time course by CFU counting on 7H10 agar plates (Methods). The assay demonstrated that the three low-level resistant mutants were also significantly tolerant to INH, with their respective time kill curves never crossing the MDK_99_ threshold, unlike the wildtype which reached MDK_99_ within 20 hours of treatment initiation (**Figure 1G**). In summary, the genotypic and phenotypic analyses revealed that while 85% of the 40 isolates may have survived low-dose INH treatment through induction of intrinsic mechanisms of phenotypic tolerance or resistance, point mutations in three genes (*ohrR*, *mfs1*, and *ntaA_5*) conferred low-level resistance, without fitness tradeoff, across 15% of the remainder isolates. Since mutations in these three genes also conferred high-level tolerance to INH, here onwards we will call these LLRT mutants.

### LLRT mutations potentiate rapid gain of high-level INH resistance with no fitness tradeoff

High-level INH^R^-conferring mutations are also associated with significant fitness tradeoff and, therefore, unlikely to survive in the absence of antibiotic within a naïve wildtype mycobacterial population. We performed the fluctuation test to investigate whether the altered fitness landscape of LLRT mutants, which we have demonstrated can co-exist in the absence of antibiotic within a naïve population, could also potentiate spontaneous gain of high-level INH^R^ (B. M. Hall et al., 2009; Jones et al., 1994). Twelve lines of the *fas1::*E177G mutant, each of the three LLRT mutants described above and wildtype Msm were inoculated at low cell densities (∼200 cells in 200 µl of 7H9 broth) and grown in the absence of antibiotic to mid-log phase. Each culture was plated on 7H10 agar with and without 50× MIC (500 µg/mL) INH and the number of INH^R^ mutant colonies and total population size were estimated with CFU counting. The FALCOR tool (B. M. Hall et al., 2009) was then used to estimate the rate of gain of INH^R^ using two independent methods, frequency and Ma-Sandri-Sarkar Maximum Likelihood Estimator (MSS-MLE) (B. M. Hall et al., 2009). The fluctuation test results showed that all three LLRT mutants acquired high-level INH^R^ at up to 6-fold higher rate, relative to the wildtype strain (**Figures 2A–B**).

**Figure 2:**
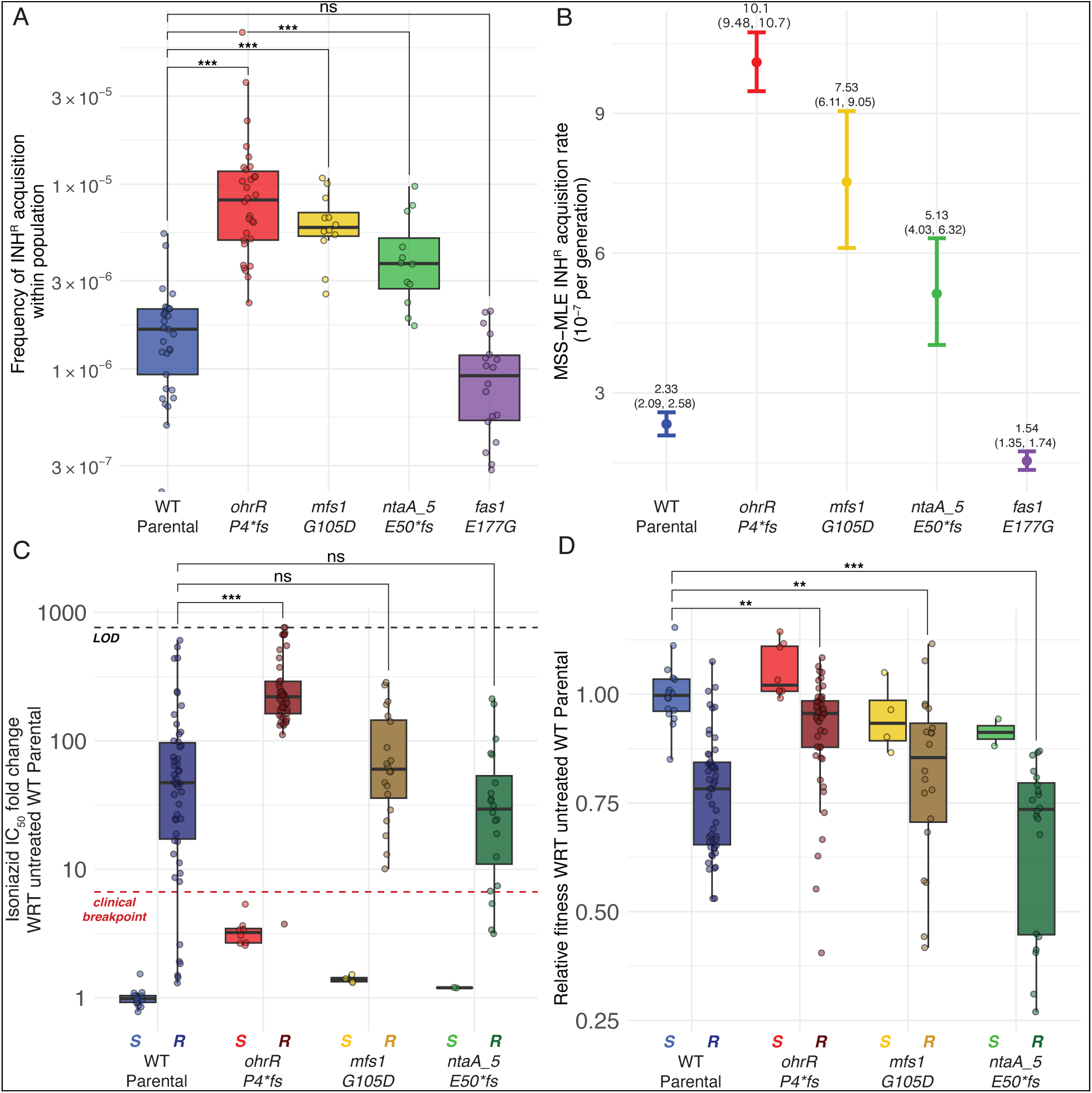
LLRT strains acquired high-level INH^R^-conferring mutations at a high rate, with minimal fitness tradeoff. Fluctuation assay results using the frequency method **(A)** or the Ma-Sandri-Sarkar Maximum Likelihood Estimator method **(B)**. Points and error bars in **(B)** represent the median and 95% confidence interval (CI), respectively. The median and upper and lower 95% CI bounds are labeled for each strain. Results are representative of two independent experiments. Fold change in INH IC_50_ **(C)** and relative fitness **(D)** of INH^R^ mutants (labeled ‘R’) and the parental strains (labeled ‘S’) from which they were derived. Relative fitness was calculated as the ratio of the AUC of each isolate with respect to the average AUC of susceptible wildtype controls after 48 hours of growth in the absence of INH. The black horizontal dashed line in **(C)** represents the upper limit of detection (LOD) for resistance, indicating ≥ 50% AUC at the maximum concentration tested relative to the strain’s untreated AUC. The red horizontal line represents the clinical breakpoint for high-level INH resistance (≥ 6.6× fold change in IC_50_). Statistical significance was evaluated using the Welch’s t-test and all p-values are Bonferroni corrected for multiple testing (\**p* < 0.05; \*\**p* < 0.01; \*\*\**p* < 0.001).

Next, we measured IC_50_ of INH in LLRT-derived INH^R^ mutants, and investigated whether the gain of resistance in these strain backgrounds was associated with fitness tradeoff in the absence of antibiotic. Dose response assays demonstrated that the level of INH^R^ acquired in LLRT strain backgrounds was comparable (*mfs1::G105D* and *ntaA_5::E50*fs*) or significantly higher (*ohrR::P4*fs*) relative to level of INH^R^ acquired in the wildtype background (**Figure 2C**). Furthermore, there was high variability in the level of INH^R^ across wildtype-derived INH^R^ mutants, suggesting a broad spectrum of evolved resistance mechanisms. By contrast, relative to INH^R^ strains derived from the wildtype background, significantly higher levels of INH^R^ were gained consistently (i.e., with low variability in IC_50_ measurements across isolates) in the *ohrR::P4*fs* background (Bonferroni corrected *p-value* = 6.87×10^−7^, **Figure 2C**). More importantly, while gain of INH^R^ was associated with significant fitness tradeoff in the wildtype, *mfs1::G105D*, and *ntaA_5::E50*fs* strain backgrounds, acquisition of INH^R^ in the *ohrR::P4*fs* strain background was associated with minimal fitness cost (**Figure 2D**). Thus, findings from the dose response and fitness assays demonstrated that LLRT mutations occur at higher frequency within a naïve population, and are enriched rapidly by brief exposure to low-dose antibiotic treatment. Further, these findings provide evidence that LLRT mutations are likely pre-resistance mutations that potentiate, with minimal fitness tradeoff, rapid gain of high-level INH^R^ in *Mycobacterium spp*.

### The *ohrR::P4*fs* mutation alleviates oxidative stress and constrains the evolutionary trajectory for acquiring high-level INH^R^

To uncover the mechanisms by which high-level of INH^R^ was acquired in the wildtype- and *ohrR::P4*fs* strain backgrounds, we sequenced the genomes of 12 small (s) and 9 large (L) INH^R^ colonies from the wildtype background and 12 small (s) and 12 large (L) INH^R^ colonies from the *ohrR::P4*fs* background. We did not see any distinguishable characteristics between small and large colonies in terms of acquired mutations, INH IC_50_, or fitness. However, while 14/21 (66%) isolates from the wildtype background had acquired nonsynonymous point mutations in *ndh* (which encodes a NADH dehydrogenase), 24/24 (100%) isolates from the *ohrR::P4*fs* background had acquired nonsynonymous frameshift mutations in the mycothiol biosynthesis (*msh*) operon genes (**Figure 3A**). By contrast, 0/21 isolates from the wildtype background harbored *msh* mutations, and 0/24 isolates from the *ohrR::P4*fs* background harbored *ndh* mutations. Interestingly, none of the isolates had gained mutations in canonical INH resistance-conferring loci aside from *ndh*. For a full list of mutations identified in the sequenced isolates, see **Table S1**. Phenotypic characterization of each of these isolates revealed that gain of high-levels of INH^R^ in the wildtype background was associated with significant fitness loss, where only 3/21 wildtype-derived INH^R^ mutants had at least 90% fitness relative to the wildtype itself in the absence of antibiotic treatment. By contrast, the majority (17/24) of the high-level INH^R^ mutants derived from the *ohrR::P4*fs* background had at least 90% fitness relative to the wildtype strain in the absence of antibiotic treatment (**Figure 3C**). These results suggest that the mechanism by which *ohrR::P4*fs* mutation buffers the fitness cost of resistance, also constrains the evolutionary trajectory (i.e., mutations in mycothiol biosynthesis genes) through which high-level INH^R^ is acquired in this specific genomic background.

**Figure 3:**
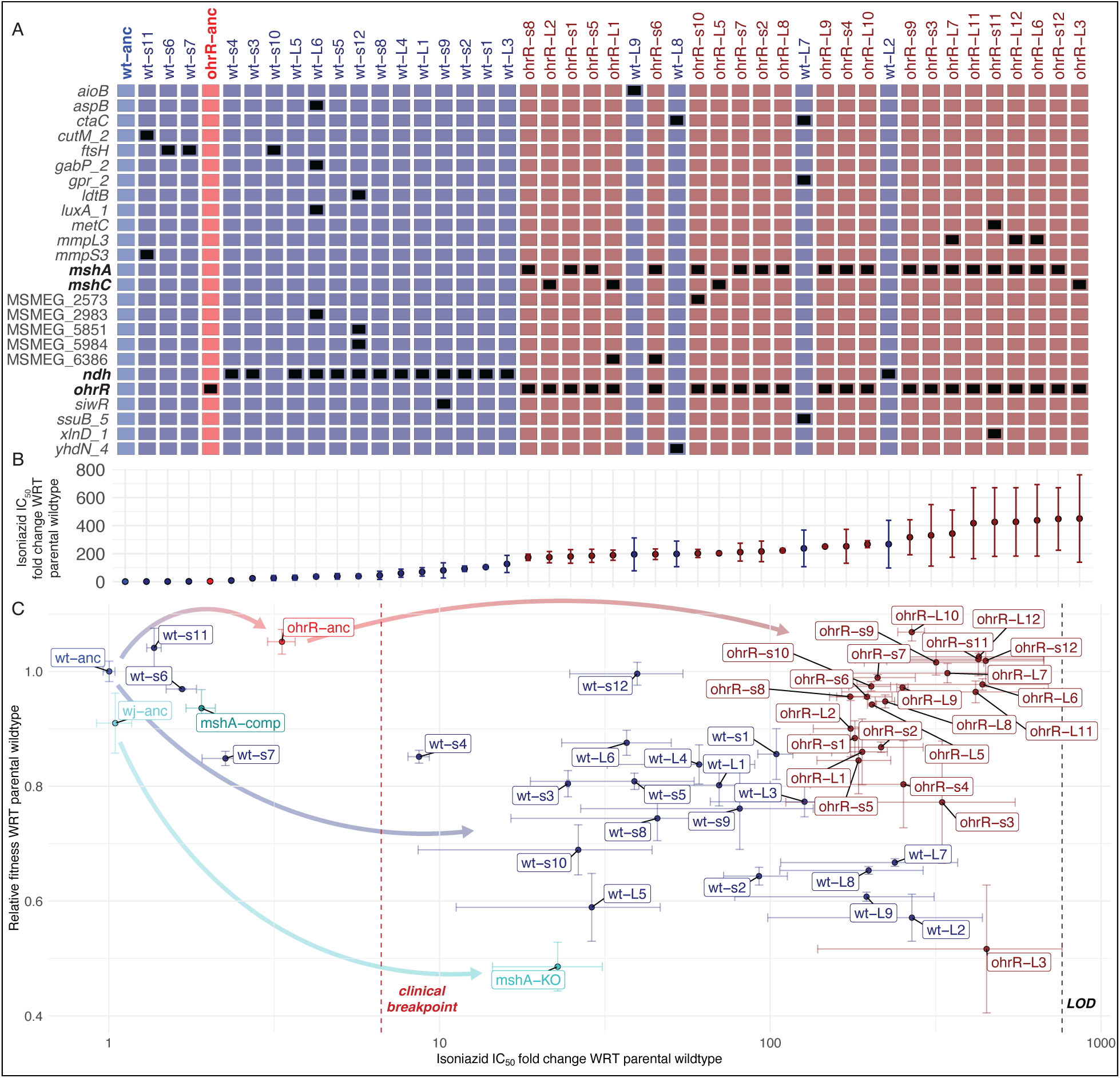
Loss-of-function mutations in *ohrR* potentiate the subsequent gain of high-level INH^R^ through a constrained evolutionary trajectory. **(A)** Grid plot of all sequenced isolates from wildtype and *ohrR::P4*fs* backgrounds, rank-ordered by their INH IC_50_. Black boxes indicate nonsynonymous mutations at specific loci (rows) in each strain (columns) identified by WGS analysis. For a full list of specific mutations, see **Table S1**. **(B)** Corresponding fold change in IC_50_ of INH of each isolate in **A. (C)** Scatter plot of relative fold changes in **A** INH IC_50_ (x-axis) and **B** fitness in the absence of antibiotic of all INH^R^ isolates derived from wildtype and *ohrR::P4*fs* backgrounds. Relative change in IC_50_ (quantified using dose-response assays) is with respect to the average IC_50_ of the ancestral wildtype strain from which the isolates were derived. Relative fitness was calculated as the ratio of the AUCs of growth curves of each isolate and the wildtype parental strain from which they were derived, after 48 hours of growth in the absence of INH. Error bars represent the standard error across 2 replicates in 2 independent experiments (n = 4).

Based on literature (Vilchèze et al., 2008) and our own findings reported here, we hypothesized that disruption of mycothiol biosynthesis by itself may confer much lower level of resistance to INH, and would carry a substantial fitness trade-off. To test this hypothesis, we performed dose response assays on an *M. smegmatis* strain with a single gene *in frame* deletion of *mshA*, and determined that while the Δ*mshA* strain had ∼22× higher IC_50_ relative to its parental strain (*p-value* = 0.0473), the *ohrR-mshA/C* double mutants generated in this study consistently showed > 200× higher IC_50_ for INH (**Figure 3C**). Furthermore, the increased INH^R^ of the Δ*mshA* single gene mutant was associated with a significant fitness tradeoff (∼48% AUC of wildtype) (**Figure 3C**), which is consistent with a prior report that MSH mutants have poor fitness due to increased oxidative stress (Vilchèze et al., 2008).

### The *ohrR::P4*fs* mutation potentiates acquisition of high level INH^R^ through the alleviation of oxidative stress

Findings from the genome re-sequencing analysis suggested that the wildtype and *ohrR::P4*fs* strains acquired high-level INH resistance through distinct mechanisms associated with mutations in the *ndh* and *mshA* genes, respectively. Disruptive mutations in *ndh* are known to confer high level INH resistance, both in Msm and in Mtb (Cao et al., 2023; Lee et al., 2001; Machado et al., 2013; Miesel et al., 1998; Rueda et al., 2015; Vilchèze et al., 2008, 2011), by increasing the NADH pool, which overrides competitive inhibition of InhA by the INH-NADH adduct (Vilchèze & Jacobs, 2014, 2019). Indeed, *ndh* mutants selected in the wildtype strain background had significantly higher NADH/NAD+ ratio with or without 1× IC_50_ INH treatment (**Figure 4A** and **Figure S2**). As expected, there was no change in NADH/NAD+ ratio in the *ohrR* or *mshA* single or double mutants, confirming a different underlying mechanism of INH^R^ in these strains.

**Figure 4:**
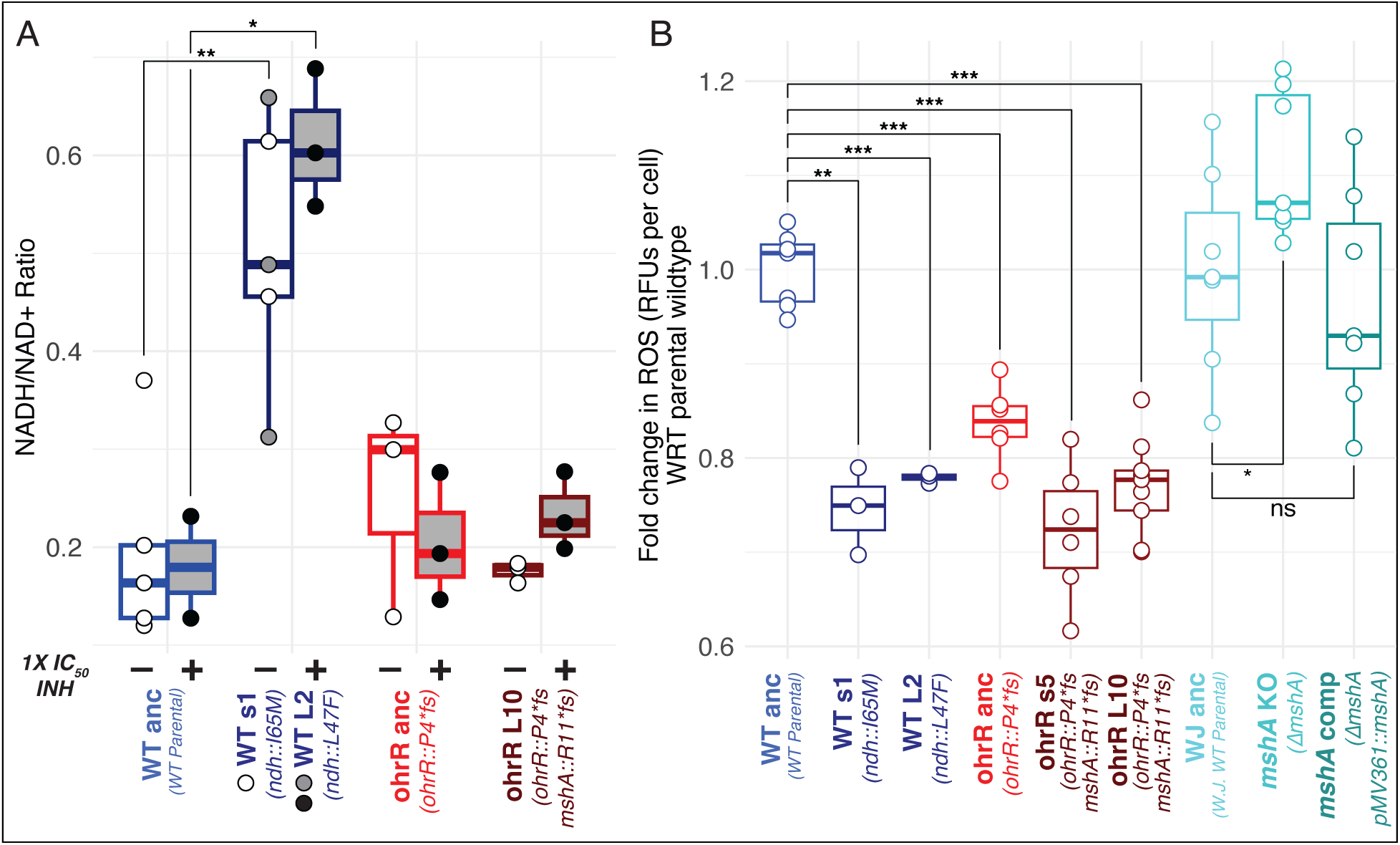
Mutations that ameliorate oxidative stress precede and potentiate the gain of subsequent high-level INH^R^ mutations. **(A)** NADH/NAD+ ratio during log phase of the wildtype parental, *ohrR::P4*fs* mutant, and INH^R^ strains evolved from each background, in the absence (indicated with “(−)”) and presence (indicated with “(+)”) of 1× IC_50_ INH. NADH/NAD+ ratios in the absence of INH were compared with respect to untreated wildtype using a Welch’s t test. Ratios in the INH-treated condition were compared similarly. **(B)** Endogenous ROS levels were measured using H2DCFDA in log phase cultures. Fold change in relative fluorescent units (RFUs) per cell was determined by comparing the RFUs per cell of each strain to the parental wildtype strain from which they were derived. Strains are labeled with the name of the background from which they were derived and their genotype relative to the wildtype parental strain. Point mutants are named by their colony size (s for small, L for large) and the isolate number. Significance was evaluated using the Welch’s t-test (\**p* < 0.05; \*\**p* < 0.01; \*\*\**p* < 0.001).

Mutations in mycothiol (MSH) biosynthesis genes, such as *mshA*, *mshB*, and *mshC*, have been associated with varying levels of INH resistance across *Mycobacterium* species, but the observed phenotypes are inconsistent and appear to depend on the broader genomic context (Rawat et al., 2007; Vilchèze et al., 2008, 2011). Studies in Msm and Mtb have reported a wide range of INH resistance levels, from low-level (∼2× MIC) to very high-level (> 100× MIC), depending on the specific mutation and strain background. In Msm, early work identified mutants deficient in MSH that were hypersensitive to rifampicin but highly resistant to INH, suggesting that MSH participates in INH activation, possibly by maintaining KatG or InhA in a reduced, reactive state (Newton et al., 1999). More specifically, *mshA* and *mshC* knockouts in Msm exhibited moderate INH resistance (4-8× MIC) alongside hypersensitivity to oxidative stress (Rawat et al., 2002; Xu et al., 2011). Similarly, *mshD::Tn5* mutants, which accumulate the thiol precursor Cys-GlcN-Ins instead of MSH, showed extreme resistance to INH (MIC > 100 µg/mL), indicating that MSH itself is required for INH activation (Koledin et al., 2002). It was also reported that transposon mutants of *mshA* (*mshA::Tn5*) exhibited very high INH resistance (> 250× MIC) (Newton et al., 2003), and that increased oxidative stress due to lack of mycothiol in these mutants was compensated by overexpression of *ohr* and ergothioneine (A. R. Singh et al., 2016; Ta et al., 2011).

Together, these findings suggest that while blocking prodrug activation in the Δ*mshA* mutant may lessen INH-induced oxidative stress (Ito et al., 1992; Lopes et al., 2020), loss of MSH increases endogenously generated oxidative stress. If confirmed, this would explain why gain of resistance to INH is less likely to occur in a wildtype background through selection of fitness-compromising loss-of-function *mshA* mutations. Independently, it has been demonstrated that knocking out *ohrR* improves the oxidative stress response (OSR) of Msm through constitutive overexpression of *ohr* (Garnica et al., 2017; Meireles et al., 2022), providing a plausible mechanism by which the *ohrR::P4*fs* pre-resistance mutation may have potentiated the gain of INH^R^ through selection of loss-of-function mutations in *mshA*. To test this hypothesis, we measured endogenous ROS levels with or without INH treatment during log-phase growth of two *ndh* mutants (*ndh::*I65M, *ndh::*L47F), two *ohrR::P4*fs - mshA::R11*fs* double mutants, Δ*mshA* and its complemented strain, as well as their parental strains. Not surprisingly, ROS levels were relatively low in the *ndh* mutants, wherein the fitness tradeoff was due to disrupted redox balance. As expected, we observed higher ROS levels in the Δ*mshA* strain, which were restored to the wildtype levels in the complemented strain. ROS levels were significantly lower in the *ohrR::P4*fs* pre-resistant mutant, and, strikingly, even lower in the INH^R^ *ohrR::P4*fs - mshA::R11*fs* double mutants (**Figure 4B**).

### Brief exposure to growth sub-inhibitory oxidative stress potentiates evolution of INH^R^

Our findings suggest that MSH-deficiency-mediated INH^R^ is unlikely to emerge in a naïve genomic background, requiring the fitness cost to be preemptively buffered by disruption of OhrR-mediated repression of Ohr. Indeed, disruption of OhrR in the *ohrR::P4*fs* strain accelerated the emergence of high-level INH^R^ by ∼6-fold through loss-of-function *mshA* mutations (**Figure 5**). Together these findings demonstrate that preemptive mitigation of oxidative stress facilitates the selection of INH^R^ mutations. Building on this, we hypothesized that the activation of the OSR through pre-exposure to low-level oxidative stress should also potentiate the evolution of INH^R^. To test this hypothesis, we performed a fluctuation assay on wildtype Msm with and without pretreatment with a growth sub-inhibitory dose (1× IC_50_; 135 ± 7.6 µM) of cumene hydroperoxide (CHx) (**Figure S5**) for 24 hours prior to plating out on 7H10 agar plates with 50× MIC INH. Indeed, pretreatment with 1× IC_50_ CHx for 24 hours increased the rate of gain of high-level INH^R^ by up to 2.7-fold relative to wildtype Msm not challenged with CHx (*p-value* = 4.51×10^-9^; **Figure 5E-F**). Taken together, our results show that activation of OSR either by genetic perturbations (e.g., disruption of *ohrR*) or pre-exposure to oxidative stress can accelerate the emergence of INH^R^ in *M. smegmatis*.

**Figure 5:**
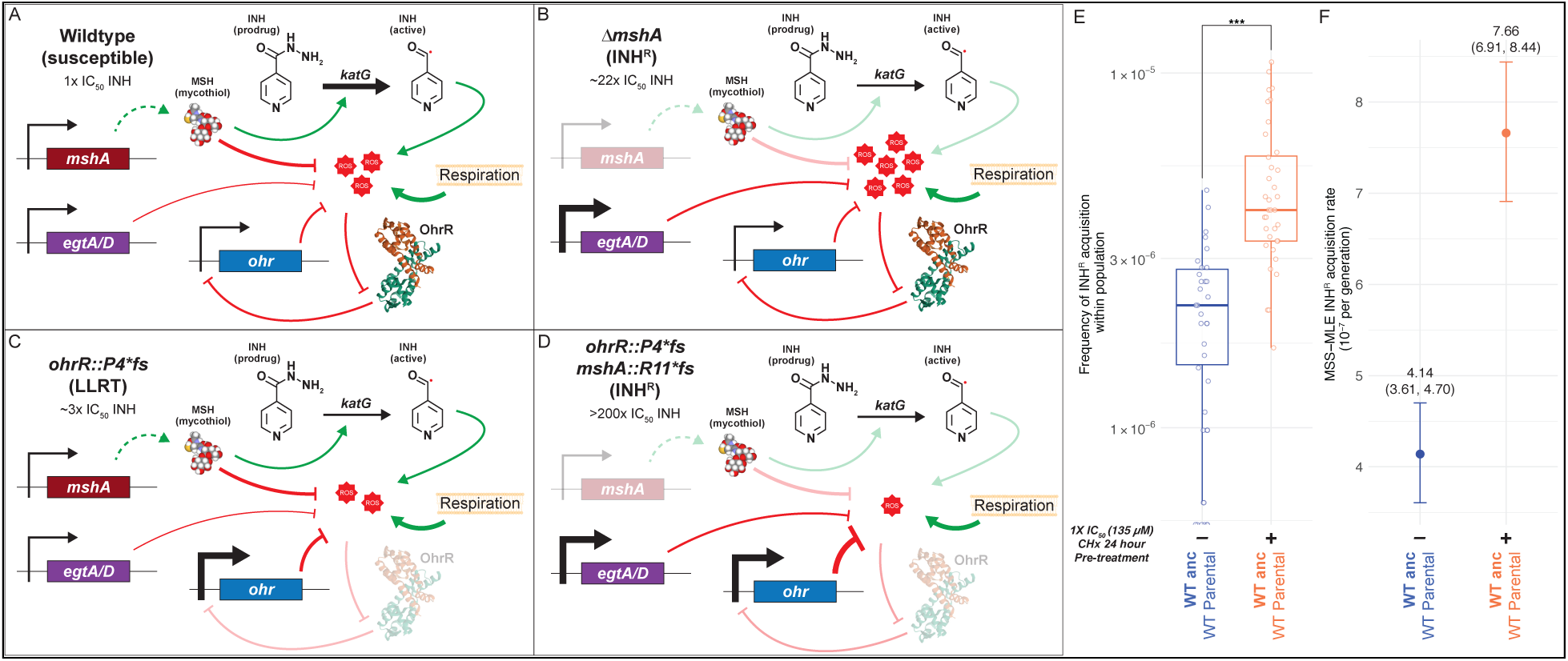
Proposed mechanism by which a loss-of-function mutation in *ohrR* supports disruptive mutations in mycothiol biosynthesis pathway genes to potentiate the evolution of INH resistance in Msm. **(A)** Genes of the *msh* operon encode biosynthesis of mycothiols (MSH). While the primary role of MSH is in mitigating oxidative stress (ROS) generated by respiration, MSH also promotes KatG-mediated activation of pro-drugs like INH (Buchmeier et al., 2003; Xu et al., 2011). Increase in ROS, such as due to INH treatment, triggers *ohr* expression by peroxide-mediated dissociation of the repressor OhrR from its promoter (Garnica et al., 2017; Meireles et al., 2022; Panmanee et al., 2006). The increased expression of Ohr helps maintain physiologically tolerable ROS levels. **(B)** While loss of MSH in the wildtype genomic background confers slightly elevated resistance to INH (∼22× IC_50_), it also results in significant increase in ROS levels, which leads to growth inhibition (loss of fitness). **(C)** Loss of function mutations in *ohrR*, such as *ohrR::P4*fs*, derepress *ohr* expression (Garnica et al., 2017; Saikolappan et al., 2014; Ta et al., 2011) resulting in significantly lower ROS levels and LLRT to INH (∼3× IC_50_). **(D).** The lower oxidative stress environment in the *ohrR::P4*fs* background supports depletion of MSH through subsequent selection of loss of function mutations in *msh* operon genes, preventing the activation of the INH prodrug and resulting in high level INH^R^ phenotype (> 200× IC_50_). The loss of the protective effect of MSH is compensated by the elevated Ohr levels (Ta et al., 2011; Vilchèze et al., 2008, 2011; Vilchèze & Jacobs, 2014). The protein model of OhrR is an orthologous structure from *Xanthamonas campestris* (PDB 2PFB) (Newberry et al., 2007). The thickness of the edges represents the model for the expected change in expression or activity of the given interaction. Fluctuation assay results to estimate rate of acquisition of INH^R^ after (+/-) pre-treatment with 1× IC_50_ CHx (135 µM), using the frequency method **(E)** or the Ma-Sandri-Sarkar Maximum Likelihood Estimator method **(F)**. Points and error bars in **(F)** represent the median and 95% CI, respectively. The median and upper and lower 95% CI bounds are labeled for each strain. Statistical significance was evaluated using the Student’s t-test (\**p* < 0.05; \*\**p* < 0.01; \*\*\**p* < 0.001).

### Bayesian analysis associates mutations in OSR genes to the potentiation and spread of strains with clinical resistance to multiple antibiotics

We investigated whether oxidative stress-ameliorating mutations may have also potentiated the emergence of INH^R^ in clinical strains of Mtb through the analysis of whole genome sequencing (WGS) data for 1,578 clinical Mtb samples from Ho Chi Minh City, Vietnam, with drug susceptibility testing (DST) results for a spectrum of frontline anti-TB antibiotics (Silcocks et al., 2023) (**Figure 6A and B**). Briefly, WGS reads were aligned to a reconstructed ancestral Mtb genome (Green et al., 2023) and analyzed with a custom variant calling pipeline adapted from (Q. Liu et al., 2022) (https://github.com/baliga-lab/bwa_pipeline) to identify 1,140,577 fixed and 194,263 unfixed (at least 10% alternative allele frequency) mutations (316,823 intergenic and 1,018,017 nonsynonymous protein-coding mutations) across 5,233 genomic loci within each sample. We then applied a Bayesian probability (BP) framework with permutation testing to the genotypic and phenotypic profiles of the Mtb samples to identify specific loci that were significantly associated with acquisition of INH resistance (**Figure 6D**) (Methods). Of the 5,233 intergenic and genic loci, mutations in 393 loci had significant BP (FDR-corrected *p-value* ≤ 0.05), of which 216 (54.96%) were protein-coding genes. As expected, protein-coding genes in loci with significant Bayes probabilities included canonical mechanisms of INH resistance, including *katG* (BP = 0.858), *inhA* (BP = 0.588), *oxyR’-ahpC* (BP = 0.667), and *katG-furA* (BP = 0.8). Significant Bayes probabilities of *pknH*-*embR* (BP = 0.714), *pyrR* (BP = 0.897), *pncA* (BP = 0.876), and *rpoB* (BP = 0.737), which are associated with canonical mechanisms of resistance to other frontline TB drugs, such as ethambutol, pyrazinamide and rifampicin, recapitulated the phenomenon of cross-resistance (Szybalski & Bryson, 1952) and evolutionary dependency (Green et al., 2023), whereby resistance to one drug (in particular INH) increases the risk of treatment failure and acquiring multi-drug resistance in TB (Gegia et al., 2017; Srinivasan et al., 2020). Interestingly, relative to their background proportion in the overall dataset (1,614/5,233, 30.84%), intergenic loci were overrepresented amongst loci with significant Bayes probabilities (177/393, 45.04%; *p-value* = 5.12×10^-10^). This finding suggested that many non-canonical regulatory mutations may have played a significant role in the emergence of INH^R^ clinical strains of Mtb.

**Figure 6:**
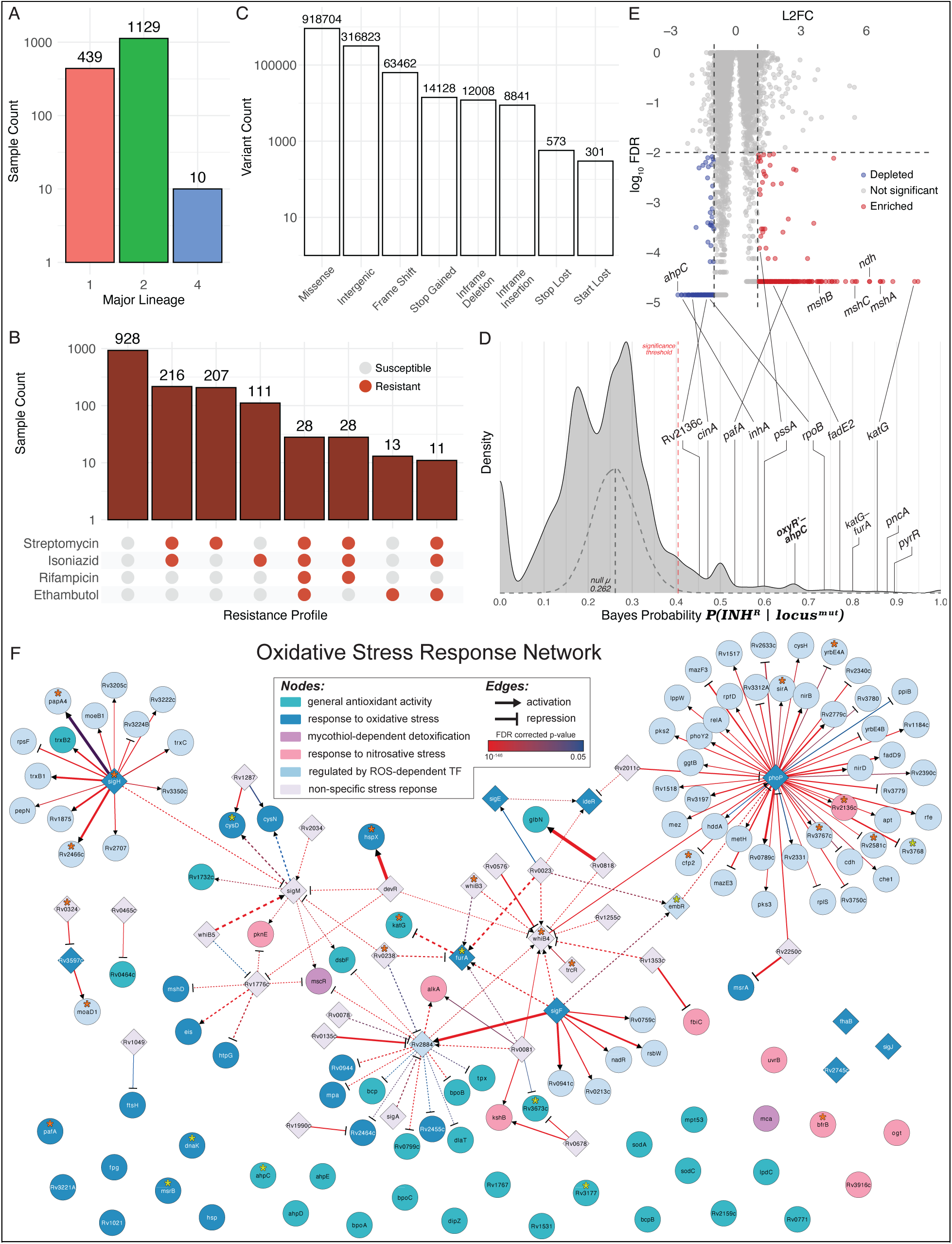
A Bayesian framework to identify genomic signatures of pre-resistance in clinical Mtb isolates. **(A)** Counts of all isolates from each major lineage, as determined by TB-Profiler. **(B)** Counts of all isolates with their corresponding susceptibility to four frontline TB drugs, as determined by drug susceptibility testing (DST) **(C)** Counts of all types of variants called across the entire dataset after removing synonymous variants from the analysis as well as variants occurring in less than 1% or more than 95% of all isolates. **(D)** Bayes probability distribution for all loci (genic and intergenic) throughout the Mtb genome for the entire dataset. The null Bayes probability distribution after > 5000 randomized permutations is overlayed as a gray dashed density plot. The significance threshold in the dashed red line represents the threshold value above which Bayes probabilities have an FDR-corrected p-value ≤ 0.05 based relative to the null distribution. Notable loci with significant Bayes probabilities are labeled on the density plot (FDR-corrected *p-value* ≤ 0.05). **(E)** Log_2_ fold change (L2FC) and false discovery rates (log_10_ FDR-corrected *p-value*) for each gene in the CRISPRi library during high-dose INH treatment (S. Li et al., 2022). This figure is a modified version of what can be found in (S. Li et al., 2022). **(D, E)** Notable loci are labeled as examples of genes with significant Bayes probability for association with INH^R^ and whose corresponding CRISPRi knockdown strains were enriched or depleted during high-dose INH treatment (S. Li et al., 2022). **(F)** Oxidative stress response network in Mtb inferred using TFOE and ChIP-seq datasets (see Methods for detailed description of how network was generated). The final network includes 159 nodes (genes) and 147 edges (TF regulatory influences). The node color represents the GO category. OSR genes without significant regulatory data were included as lone nodes at the bottom of the network. The shape of the edge end represents repression (bar) or activation (arrow). Bold edges represent interactions with evidence of TF influence (TFOE) and binding (ChIP-seq). The dashed edges represent interactions with only TFOE evidence. The color of the edge represents the TFOE FDR-corrected p-value, and the width of the edge represents the TFOE log_2_ fold change in expression. Loci with significant BP for potentiating INH resistance are notated with an asterisk, with the color depending on if the significant probability corresponds to the gene itself (orange asterisk) or the upstream, intergenic region (yellow asterisk).

Using results from a genome-wide fitness screen, we then discovered that the 207/216 genes included in the screen with significant Bayes probabilities were significantly overrepresented (hypergeometric test *p-value* = 0.0261) among CRISPRi knockdown strains that were depleted (19/207 genes) or enriched (9/207 genes, FDR ≤ 0.01, absolute value log_2_ fold change ≥ 1) after 5 days of high-dose INH treatment (S. Li et al., 2022) (**Figure 6E**). Given the interesting finding that the Bayesian analysis had uncovered pre-resistance mutations that may have potentiated multidrug resistance, we also analyzed the CRISPRi screening results for evidence supporting the functional association of high BP genes with resistance to other antibiotics. This analysis demonstrated that although the Bayesian analysis was performed in the context of INH^R^, CRISPRi strains targeting high BP genes were significantly enriched or depleted by treatment with multiple other antibiotics, including rifampicin, ethambutol, clarithromycin, streptomycin, and vancomycin (**Table 2**).

**Table 2.**
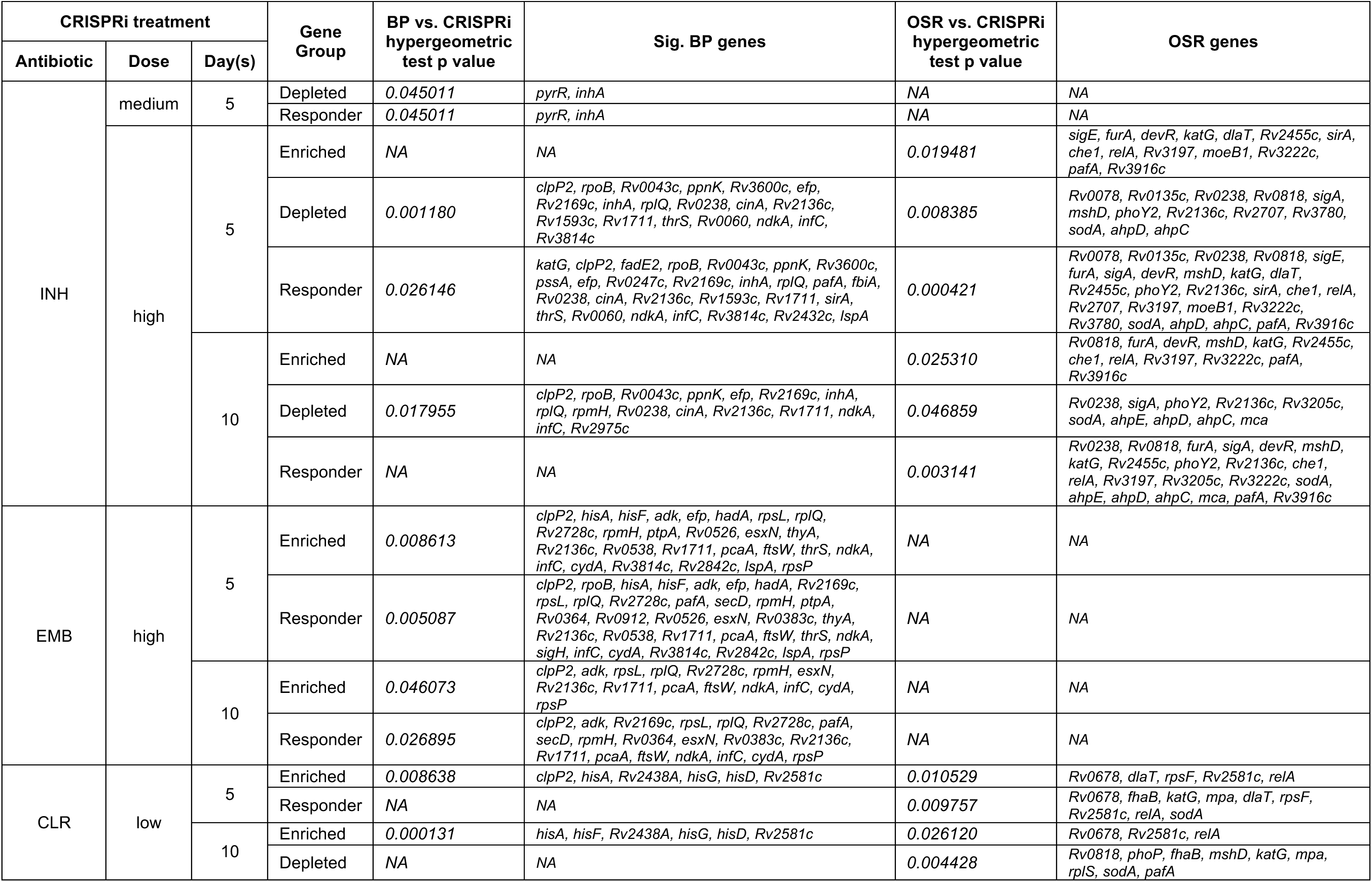
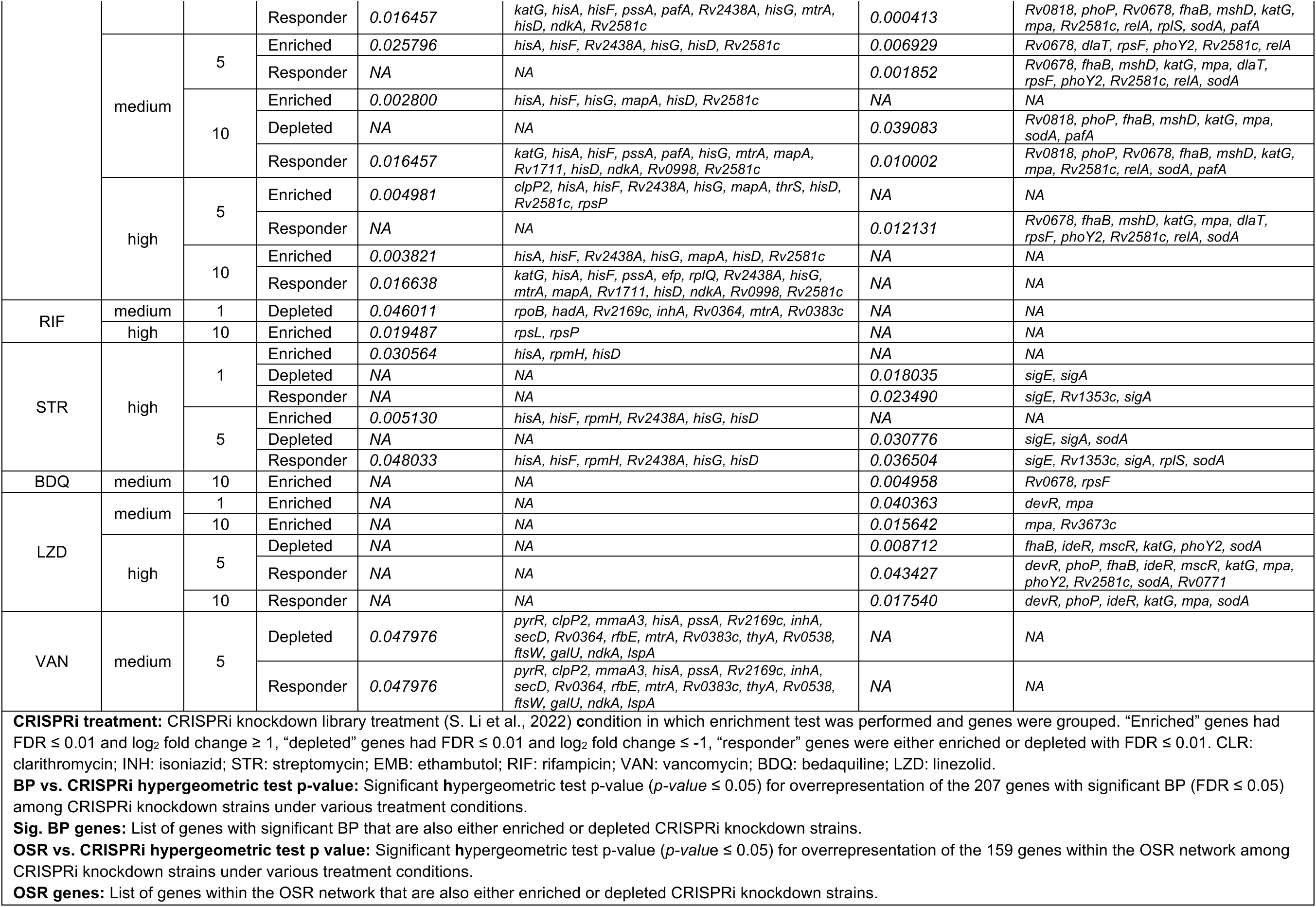
Overrepresentation of genes with significant BP or OSR network genes among CRISPRi knockdown strains during antibiotic treatment.

Notably, CRISPRi knockdown strains of *mshA*, *mshB*, and *mshC* were among the most enriched during treatment with INH, providing additional evidence for fitness advantage of disrupting mycothiol biosynthesis in Mtb during INH treatment. The high Bayes probability of the *oxyR’-ahpC* intergenic region as well as the finding that the CRISPRi knockdown strain of *ahpC* was among the most depleted after 5 days of high-dose INH treatment, underscored the importance of oxidative stress mitigation of AhpC function, the Mtb ortholog of Ohr, and its regulation in the emergence of INH resistance. Given that Mtb routinely experiences oxidative stress inside the host during infection, we further explored whether mutations in genes generally involved in the mitigation of oxidative stress may have potentiated the emergence of INH resistance. In order to do this, we first compiled a list of 68 genes associated with mitigation of oxidative stress in Mtb by leveraging assigned gene ontology (GO) terms (antioxidant activity, GO:0016209; cellular oxidant detoxification, GO:0098869; removal of superoxide radicals, GO:0019430; mycothiol-dependent detoxification, GO:0010127; response to oxidative stress, GO:0006979; response to nitrosative stress, GO:0051409). Then, we constructed a regulatory network for these genes by leveraging two independent datasets: the transcription factor over-expression (TFOE) dataset that catalogs sets of genes that are differentially expressed as a consequence of overexpressing any of 209 TFs in the Mtb genome (Rustad et al., 2014) and experimentally-mapped binding sites for 143 TFs through chromatin immunoprecipitation sequencing (ChIP-seq) (Minch et al., 2015) (Turkarslan et al., 2015) (Methods). The final 159 gene OSR network constructed using GO, ChIP-seq and TFOE, includes all known GO-annotated OSR genes (58 genes and 10 TF regulators), their own 27 TFs, as well an additional 64 downstream target genes (**Figure 6F**). Strikingly, the final OSR network of Mtb (122 genes regulated by 37 TFs) was enriched for 19 genes and 9 intergenic regions with significant BP for potentiating INH resistance (hypergeometric test *p-value* = 1.09×10^-7^). Many genes within the OSR network that also have significant BP have been previously implicated in mono- or multi-drug resistance, including Rv2136c (Ehrt & Schnappinger, 2009), *pknH*-*embR* (Sharma et al., 2006; Sun et al., 2023; Belanger et al., 1996), *whiB3* (A. Singh et al., 2007), *katG*-*furA* (Ando et al., 2011; Baker et al., 2005; Milano et al., 2001; Pym et al., 2001), Rv0238 (Ortega Ugalde et al., 2019), and *sigH* (Cioetto-Mazzabò et al., 2023; A. K. Singh & Singh, 2009) (**Table S2**). Upon further examination of overlap with additional CRISPRi screens, we discovered that the OSR network genes were also significantly associated with Mtb survival in the presence of multiple other antibiotics, including bedaquiline, clarithromycin, linezolid, and streptomycin (**Table 2**). Thus, findings from our studies on Msm, the Bayesian analysis of clinical strains of Mtb, CRISPRi knockdown screens (S. Li et al., 2022), and the analysis of the OSR network of Mtb, together demonstrated that mutations that mitigate oxidative stress may have played an important role in the emergence of multi-drug resistance in Mtb.

## Discussion

The findings presented in this study reveal that oxidative stress management and exposure to low-dose antibiotic may play a critical and previously underappreciated role in the evolution of antibiotic resistance in *Mycobacterium tuberculosis*. The data support a model wherein LLRT mutations that enhance oxidative stress resilience, such as those affecting the OxyR-AhpC regulatory axis, create a permissive genomic background that accelerates the acquisition of high-level drug resistance without incurring prohibitive fitness costs.

### OxyR-deficient mutants are oxidative stress pre-adapted populations that potentiate rapid emergence of INH^R^ strains in clinical Mtb

The Bayesian analysis of clinical Mtb samples from Vietnam recapitulated an established association between mutations in the *oxyR’-ahpC* intergenic region and INH resistance. This association is supported by the well-established role of OxyR’ (the Mtb ortholog of OhrR) as a negative regulator of *ahpC*, which encodes alkyl hydroperoxide reductase – a key enzyme in detoxifying reactive oxygen and nitrogen species (Sherman et al., 1999; Heym et al., 1997; Kelley et al., 1997; Wilson & Collins, 1996), associated with increased resistance to oxidative stress (Dhandayuthapani et al., 1996). Furthermore, both depletion of the CRISPRi knockdown strain of *ahpC* in the genome-wide fitness screen in high-dose INH (S. Li et al., 2022) as well as the complete absence of loss-of-function mutations in *ahpC* across clinically resistant Mtb strains (Comas et al., 2013) support the essentiality of *ahpC* for INH^R^ TB infections. While previously it was believed that constitutive AhpC overexpression, through loss-of-function or regulatory mutations in *oxyR’*, was a compensatory mechanism (Andersson & Hughes, 2010) that appeared later to support INH resistance (primarily due to loss of function mutations in *katG*) (Sherman et al., 1999), our findings suggest that *ahpC* overexpression mutations likely appeared first and potentiated the selection of high level INH resistance mutations. In this regard, the hypothesis that OxyR-deficient mutants may naturally exist within naïve Mtb populations (Dhandayuthapani et al., 1996) is strongly supported by the biology of tuberculosis. Mtb resides primarily within host macrophages, where it is subjected to sustained oxidative and nitrosative stress as part of the innate immune response (Ehrt & Schnappinger, 2009; Voskuil et al., 2011). In this hostile environment, bacteria with enhanced oxidative stress defenses, such as those with derepressed *ahpC* expression, would be expected to have a selective advantage even in the absence of antibiotic pressure. Thus, *oxyR’-ahpC* promoter mutations are likely "pre-resistance" mutations (Levin-Reisman et al., 2017) that incur minimal fitness cost under baseline conditions but prime Mtb populations for rapid acquisition of INH^R^ with subsequent selective pressures during INH treatment.

### Oxidative Stress Exposure Accelerates Resistance Evolution

The significant enrichment of oxidative stress-related loci among genes with high BP for association with INH resistance, further underscores the point that constitutive activation of oxidative stress defenses is not merely correlated with resistance but mechanistically linked to it. Specifically, this finding suggests that mutations in multiple genes associated with management of oxidative stress may have acted as pre-resistance mutations that led to the emergence and spread of INH^R^ strains in Vietnam. Mathematical modeling of antibiotic tolerance and resistance emergence supports this interpretation (Levin-Reisman et al., 2017). Specifically, through model simulations Levin-Reisman et al. showed that tolerance mutations – those that transiently allow survival under antibiotic stress without conferring high-level resistance – can arise more frequently and in a broader range of genes than direct resistance mutations, effectively serving as evolutionary stepping stones toward resistance. Consistent with this theory, our findings show that, with over 150 genes, Mtb’s OSR network provides a much larger target for acquiring LLRT mutations without major fitness costs compared to the few genes involved in canonical high-level drug resistance. In fact, the high ROS within the host microenvironment likely favors selection of these oxidative stress ameliorating LLRT mutations creating fertile evolutionary ground for the subsequent selection of stable, high-level resistance. Additional support for this theory comes from our tolerance assays, which showed that the slow rate of killing of all Msm LLRT mutants by 100× MIC INH was immediately followed by regrowth within 60 hours of treatment initiation, likely due to the rapid emergence of high-level INH^R^ mutants. Together, these findings support a model in which oxidative stress first selects for LLRT mutations within the OSR network, thereby accelerating the evolutionary trajectory toward fixed, high-level drug resistance. The experimental demonstration that brief exposure to sub-inhibitory oxidative stress (via low-dose cumene hydroperoxide treatment) nearly tripled the rate of evolution to high-level INH resistance in Msm further provides compelling evidence that the oxidative environment within the host may also actively promote drug resistance evolution in Mtb.

This finding has important implications for understanding the epidemiology of drug-resistant tuberculosis. Patients with active TB experience chronic inflammation, and the granulomas that characterize tuberculosis pathology are sites of intense oxidative stress (Amaral et al., 2021; Palanisamy et al., 2011; C.-T. Yang et al., 2012). Our data suggest that this hostile microenvironment may inadvertently select for or enrich mutants with enhanced oxidative stress defenses, such as those with *oxyR’* mutations, thereby creating a reservoir of pre-resistant bacteria primed to rapidly acquire full resistance upon antibiotic exposure. Furthermore, recent studies have demonstrated that heterogeneity in drug penetration within granulomas creates microenvironments where bacteria experience sub-inhibitory antibiotic concentrations (Cicchese et al., 2020), which may also enrich for LLRT mutants as we have shown, further compounding the selective pressure for pre-resistance.

Our finding that *ohrR* loss-of-function LLRT mutants in *M. smegmatis* acquire high-level INH resistance at > 6-fold increased rate, primarily through subsequent mutations in mycothiol biosynthesis genes, provides a mechanistic explanation for this phenomenon for one such pre-resistance mutation in Msm. Loss of mycothiol, a key antioxidant in mycobacteria, is typically detrimental due to increased sensitivity to endogenously generated oxidative stress (Buchmeier et al., 2003; Rawat et al., 2002), and is particularly problematic for slow growing mycobacteria, such as Mtb, which accumulate more oxidative damage per cell division (Bhaskar et al., 2014). However, when alternate mechanisms of defense against oxidative stress are constitutively upregulated (as in *ohrR* mutants), the fitness cost of *mshA* mutations is buffered, allowing these otherwise deleterious mutations to be retained. Importantly, *mshA* mutations block the activation of INH by KatG, thereby conferring resistance through reduced drug activation rather than target modification (Vilchèze et al., 2008; Xu et al., 2011). This two-step evolutionary trajectory, wherein oxidative stress-mitigating mutations precede and enable the selection of resistance-conferring mutations, may explain the rapid emergence of high-level resistance observed clinically.

### Clinical and Therapeutic Implications

The findings presented here have several important implications for tuberculosis treatment and drug development. First, they suggest that monitoring for mutations in OSR genes, particularly *oxyR’-ahpC*, may serve as an early warning for populations at elevated risk of acquiring drug resistance. Mutations in these loci may represent actionable biomarkers that warrant more aggressive treatment regimens or closer monitoring for resistance emergence. Second, therapeutic strategies aimed at disrupting oxidative stress buffering systems may synergize with existing antibiotics. For instance, compounds that inhibit AhpC or other oxidative defense enzymes (Koshkin et al., 2004; Ren et al., 2020) could potentially reverse the permissive environment created by pre-resistance mutations, thereby reducing the likelihood of resistance evolution. Third, these findings underscore the importance of ensuring adequate drug penetration and compliance in TB treatment. The prolonged periods of sub-inhibitory drug exposure that can occur due to granuloma heterogeneity or treatment non-compliance create conditions that favor both the enrichment of pre-resistant mutants and the subsequent selection of high-level resistance (Andini & Nash, 2006; Cicchese et al., 2020; Pablos-Méndez et al., 1997; Sarathy et al., 2016). Thus, our findings provide further support for the importance of strategies to improve drug delivery to granulomas or to reduce treatment duration without compromising efficacy to help mitigate the risk of emergence of drug resistance.

### Limitations and Future Directions

Several limitations should be noted. The dataset of 1,578 Mtb isolates, while valuable for linking genotype to experimentally determined phenotype, represents a single geographic region and does not fully capture global diversity. Additionally, the Bayesian framework employed here cannot definitively distinguish causal mutations from those linked by genetic hitchhiking, nor can it fully resolve the temporal sequence of mutational events. Longitudinal studies tracking the emergence of oxidative stress-related mutations in patients over the course of treatment would be valuable for validating these findings.

Our findings also suggest that high BP genes, especially those involved in the OSR network, may have also potentiated gain of resistance to other anti-TB drugs. Future work should explore this finding further, specifically, whether similar pre-resistance mechanisms operate for other anti-TB drugs and whether oxidative stress-mitigating mutations also potentiate resistance to drugs beyond INH. Given that many drugs act through generation of oxidative stress (Dwyer et al., 2014), the principles identified here may have broad applicability with regard to understanding how including such drugs in combination regimen may potentiate the evolution of multidrug resistance. Conversely, the development of adjunct therapies targeting oxidative stress defenses, combined with traditional antibiotics, represents a promising avenue for clinical intervention.

## Conclusion

This study demonstrates that oxidative stress management is a critical determinant of antibiotic resistance evolution in *M. tuberculosis*. Mutations that enhance oxidative stress resilience, likely pre-existing in clinical populations due to selection by the host immune environment, create a genomic background that accelerates the acquisition of high-level drug resistance without fitness trade-offs. These findings challenge the traditional view of resistance evolution as driven primarily by antibiotic exposure and highlight the importance of considering the host environment in shaping emergence of multidrug resistance. Ultimately, targeting oxidative stress pathways in the pathogen may represent a novel strategy for preventing or delaying the emergence of drug-resistant tuberculosis.

## Materials and Methods

### Bacterial strains and growth conditions

All *M. smegmatis* strains were derived from the parental mc^2^155 strain, provided to us by the Bhatt lab at Stanford University. Additional strains W.J. Wildtype (mc^2^155 parental strain), *mshA KO* (in-frame deletion of *mshA* via phage transduction), and *mshA comp.* (*mshA KO* complemented with episomal plasmid expression of functional *mshA*) were kindly provided by the Jacobs lab at the Albert Einstein College of Medicine. *M. smegmatis* was grown at 37°C in Middlebrook 7H9 broth or 7H10 plates supplemented with 0.2% glycerol, 0.05% Tween-80 (liquid media), and 10% OADC, with aeration. Media for all species of bacteria was free of antibiotics unless otherwise noted.

### Enrichment and selection of *M. smegmatis* LLRT mutants

Biological replicates of Msm were picked off of agar and inoculated into ∼6 mL of media and grown until the cultures reached mid-log phase (OD_600_ 0.6-1.0). Then, the OD_600_ of each replicate culture was normalized to 0.2 into 6 mL media + 2× IC_50_ antibiotic (INH). Cultures were then incubated for 16 hours to allow pre-resistant mutants to be enriched while also preventing the culture from undergoing an entire doubling, on average. Cultures were then plated on agar with or without 2× IC_50_ antibiotic and then positioned on top of an image scanner placed inside an incubator. ScanLag was performed by capturing an image of sets of plates every hour for ∼72 hours (Levin-Reisman et al., 2010). Images were then compiled and colonies were detected and their growth metrics were inferred using the ScanLag software, available at https://github.com/baliga-lab/Scanlag.git. Dimensionality reduction with PCA was performed on the scaled measurements of time of appearance, growth rate, and maximum colony size and unsupervised clustering using k-means with k = 3 was performed to group phenotypically distinct colonies together. K-means clustering of the PCA using k = 3 was determined based on the within sum of square distance from each cluster center. Representative colonies from each cluster were then picked and assayed.

### Dose response assay for fitness and IC_50_ determination

Dose response assays were performed in 384-well black-wall plates (Greiner Microplate, 384 Well, Cat # 781097) and growth within each well was recorded as the OD_600_ with a plate reader spectrophotometer. Wells with cultures were seeded at an initial OD_600_ = 0.01 and blank wells were filled with sterile media. A log_2_-fold dilution series of antibiotic concentrations ranging from 0× MIC to roughly 4-10× MIC was aliquoted out to their corresponding wells. Shortly after untreated cultures reached stationary phase, the experiment was stopped and the data was processed in R (version 4.1.1) (R Core Team, 2024). Briefly, the mean of each blank OD_600_ at each time point was subtracted from the OD_600_ of the wells containing cultures. The area under the curve, lag phase, and additional growth metrics were inferred with GrowthCurver (version 0.3.1) (Sprouffske & Wagner, 2016). Relative fitness was calculated as the ratio of the area under the growth curve (AUC) of the strain being analyzed during uninhibited growth (no antibiotic) relative to the average AUC of all untreated or wildtype control strains. To determine IC_50_ values, a log-logistic model was fit to the dose response data using the drc package in R (version 3.0-1) (Ritz et al., 2015).

### Tolerance assays

Twelve biological replicates of each tested strain were inoculated in fresh 7H9 media and grown overnight to mid-log phase (OD_600_ of ∼0.5) and diluted to OD_600_ of 0.05, corresponding to ∼10^7^ CFU/mL in 3 mL of fresh 7H9. Isoniazid was added to each sample to a final concentration of 1 mg/mL (100× wildtype MIC). Cultures were incubated at 37°C shaking at 225 RPM and subsampled at the following time points: 0, 6, 24, 30, 48, and 72 hours. Aliquots were serially diluted 1:5 in fresh antibiotic-free 7H9 and a spotting assay was performed by plating out 4 µL of each dilution on 7H10 agar in an OmniTray (Thermo Sci Cat #140156). Agar plates were then incubated at 37°C for 2-3 days until colonies appeared, at which point they were imaged and colonies were counted for CFU determination.

### Fluctuation assays

12 biological replicates of each strain used in these experiments were inoculated and grown to mid-log phase (OD_600_ of ∼0.5-0.8) in the absence of antibiotics. OD_600_ of each culture was measured and normalized into 200 µL of media in a 96-well plate such that each well contained no more than ∼200 cells. The plate was incubated with shaking until cultures reached late-log phase (OD_600_ of ∼1.0). Growth was stopped before cultures entered stationary phase to prevent the emergence of mutations that arise as a result of nutrient starvation. 40 µl of culture was serially diluted and 4 µl of each dilution was spotted on agar to determine the CFU/mL (population size). 100 µl of remaining culture in each replicate was plated on agar with 50× MIC INH. After colonies appeared, they were counted and mutation rates were determined using the web-based FALCOR tool (https://lianglab.brocku.ca/FALCOR/) using the frequency method and the Ma-Sandri-Sarkar Maximum Likelihood Estimator (MSS-MLE) method (B. M. Hall et al., 2009). For fluctuation assays including pre-treatment with 135 µM cumene hydroperoxide (Thermo Fisher Cat #L06866), 1× IC_50_ (as determined by dose response assay) was added to log-phase cells in 7H9 minimal media (7H9 + 0.2% glycerol, 0.05% Tween-80) to prevent degradation of the cumene hydroperoxide by catalase in the media for 24 hours before cultures were plated on agar with and without 50× MIC INH for counting CFUs and mutant colonies exactly as described above. Mutation rates were adjusted for the difference in the number of generations the treated and untreated cells underwent before plating.

### NADH/NAD+ ratio determination

NADH and NAD+ concentrations were measured using the Enzychrom NAD/NADH assay kit (Bioassay Systems), following the manufacturer’s instructions (and adding a bead-beating step of 15 minutes before heating at 60°C for 5 minutes to lyse the bacterial cells). The equivalent of 1 mL of OD_600_ of 1.0 bacterial cells (∼10^8^ CFUs) were used to measure NADH and NAD+ concentrations. To capture bacterial cells in the same growth phase in the treated and untreated conditions, these experiments were performed separately on different days. In the absence of isoniazid, log-phase cultures were directly OD normalized and processed. For the treated condition, log-phase cultures were normalized to OD_600_ of 0.95 and 1× IC_50_ isoniazid (3.0 µg/mL for WT Parental, 270 µg/mL for WT s1 (*ndh::I65M*), 700 µg/mL for WT L2 (*ndh::L47F*), 9.4 µg/mL for ohrR anc (*ohrR::P4*fs*), and 770 µg/mL for ohrR L10 (*ohrR::P4*fs*, *mshA::R11*fs*) was added and cultures were incubated at 37°C shaking at 225 RPM for 2 hours before NADH and NAD+ quantification. 1× IC_50_ isoniazid was used to minimize the impact of fitness differences between the strains in the measured NADH/NAD+ ratio. Final results included at least two replicates per strain.

### ROS assay

At least three biological replicates of each indicated strain were grown to mid-log phase (OD_600_ of 0.5-1.0) in 7H9 minimal media (7H9 + 0.2% glycerol, 0.05% Tween-80) and then normalized to 8 × 10^6^ cells/mL. 200 µl replicates were added to black-walled 96-well plates (Thermo Sci Cat #237107). The fluorescent dye 2’,7’-dichlorodihydrofluorescein diacetate (H2DCFDA, Ex/Em: ∼492–495/517–527 nm, Thermo Fisher Cat #D399) was added to OD normalized cultures of all indicated strains to a final concentration of 10 µg/mL, plates were sealed with foil sealing tape, and incubated at 37°C for 1 hour. RFUs were measured in a BioTek Synergy (H4 Hybrid Reader) fluorescence plate reader at the indicated excitation and emission wavelengths and fold change was calculated for each strain with respect to the wildtype parental strain in which each strain was derived.

### Whole genome sequencing

Mid- to late-log phase cultures were harvested by centrifugation and genomic DNA was extracted and purified with the Qiagen DNeasy UltraClean Microbial Kit (Cat #12224). Libraries for sequencing were prepared with the Nextera XT DNA library preparation kit (Illumina, San Diego, CA) and pooled for paired-end sequencing on a NextSeq instrument.

### Identification of mutations in evolved and wildtype Msm strains

Raw fastq files generated from Illumina sequencing were processed and mutations were called using a custom variant calling pipeline (https://github.com/baliga-lab/bwa_pipeline). Variant calling was performed with GATK (version 4.3.0.0) (DePristo et al., 2011), Varscan (version 2.4.3) (Koboldt et al., 2012) and bcftools from Samtools package (version 1.6) (H. Li et al., 2009). Variants identified by each caller were collated and filtered for variant frequency ≥10%. Variants called by at least two algorithms were included for further analysis, including annotation using SnpEff (version 5.1d) (Cingolani et al., 2012). The reference genome used for *M. smegmatis* was mc^2^155: NC_008596.

### Identification of mutations in public Mtb WGS data

We harnessed whole genome sequencing data from the Sequence Read Archive of 1,578 clinical Mtb samples from Ho Chi Minh City, Vietnam (Silcocks et al., 2023). The function *fasterq-dump* of the SRA toolkit (https://www.ncbi.nlm.nih.gov/books/NBK242621/) was used to download the raw *fastq* files and convert them to paired-end *fastq* files (“Download Guide,” 2016). Reads were aligned to the reconstructed ancestral Mtb genome (Green et al., 2023) and mutations were called using the same custom variant calling pipeline as above, adapted from (Q. Liu et al., 2022), designed to capture fixed (≥ 90% alternative allele frequency) and unfixed (≥ 10% and ≤ 90% alternative allele frequency) intergenic and protein-coding mutations within each sample. As done in (Q. Liu et al., 2022), samples with an average sequencing depth over 20× and a mapping rate of over 90% were used for downstream analyses. Additionally, we excluded synonymous variants and variants located in regions that are difficult to characterize with short-read sequencing, including repetitive regions of the genome such as PPE/PE-PGRS family genes, pro-phage genes, insertions or mobile genetic elements (Marin et al., 2022). Computationally predicted lineage subtypes for each sample were identified using the TB-Profiler tool (version 6.3.0) (Phelan et al., 2019).

### Bayesian analysis of Mtb metagenomes

After calling variants and collecting drug susceptibility profiles of each Mtb genome included in the dataset, an occurrence matrix was generated for all loci with at least one mutation in at least 1% of samples to associate mutation occurrence with drug susceptibility. To do this, we took the genotypic and phenotypic profiles of each sample, and used a Bayesian framework to calculate the conditional probability of an isolate being INH drug resistant given a mutation in any locus in the genome. We applied the classic Bayes Theorem:

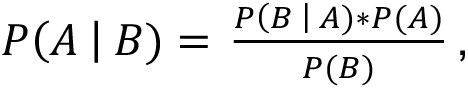

where *P(A)* is the probability of being resistant to a particular antibiotic (i.e., proportion of all samples resistant to INH), *P(B)* is the proportion of samples with a mutation in the locus of interest, and *P(B|A)* is the proportion of samples predicted to be drug resistant that also harbor a mutation in the locus of interest. Contextualized, Bayes Theorem can we written as:

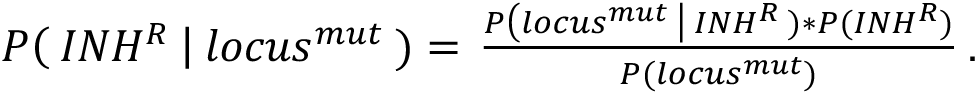

This expression can then be expanded to show how each component is calculated, enabling us to simplify the equation:

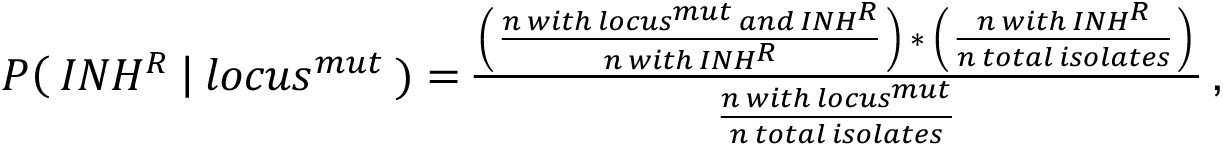

where *n* is the number of samples with WGS that correspond to that given category (i.e. *‘n total isolates’* is the total number of samples in the dataset, *‘n with locus^mut^’* is the total number of samples with at least one mutation in the specific locus). This expression can be simplified by canceling out like terms to arrive at the final equation used to calculate Bayes Probabilities:

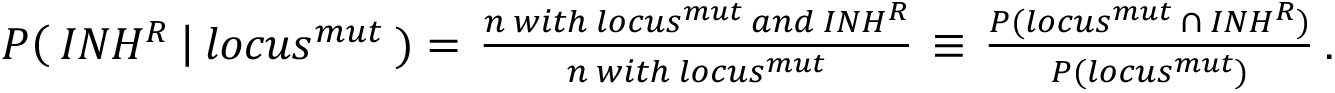

We performed the conditional probability calculation for each genic and intergenic locus in the Mtb genome and then assigned p-values to each probability by comparing the value to the distribution of Bayesian probabilities for all loci across > 5000 iterations of randomizing the mutation profiles of all samples in the dataset. p-values were then FDR corrected for multiple testing and a significant Bayes probability was determined to be any loci with an FDR-corrected *p-value* ≤ 0.05.

### Development of oxidative stress response network in Mtb

A list of genes was compiled based on gene ontology for functions related to the following GO categories: antioxidant activity, GO:0016209; cellular oxidant detoxification, GO:0098869; removal of superoxide radicals, GO:0019430; mycothiol-dependent detoxification, GO:0010127; response to oxidative stress, GO:0006979; response to nitrosative stress, GO:0051409. We assembled a draft network by identifying experimentally predicted interactions and binding sites from transcription factor over-expression (TFOE) (Rustad et al., 2014) and chromatin immunoprecipitation sequencing (ChIP-seq) (Minch et al., 2015) datasets, which were further summarized in (Turkarslan et al., 2015). The TFOE dataset was first filtered to only include influences with an FDR-corrected *p-value* ≤ 0.05. Then, each of the 206 TFs included in the TFOE dataset were assessed by hypergeometric test to identify regulons enriched for genes associated with ROS/RNS stress. Regulons enriched for oxidative stress-related functions (hypergeometric test *p-value* ≤ 0.05) were retained (5 TFs, 493 nodes, 531 edges). The network was then filtered to only include genes from the GO analysis (7 TFs, 28 nodes, 28 edges). The predicted transcriptional regulators (based on the analyzed experimental data) of the TFs with enriched regulons (25 TFs, 26 nodes, 40 edges) were then also added. Next, to include high-confidence regulatory relationships for genes that were filtered out during the enrichment analysis, TF-target interactions for TFs or targets in the GO gene list with significant TFOE influences and ChIP-seq binding evidence (18 TFs, 97 nodes, 89 edges) were also included in the network. Lastly, genes among the 68 identified from GO analysis that were left out of the network due to insufficient regulatory data (28 genes, 3 TFs) were manually added back to the network as lone nodes. The final network contained 37 TFs, 159 nodes and 147 edges and was visualized with Cytoscape v3.10.3 (Shannon et al., 2003) and analyzed with the NetworkAnalyzer (Assenov et al., 2008) tool to identify node outdegrees.

## Code and data availability

All custom code used to generate results and figures reported in this manuscript can be found at https://github.com/evanpepper/Mycobacterium_PreR along with experimental data and supplementary files. Reviewers are encouraged to clone the repository and run the code (R Markdown) to reproduce the reported results. Susceptibility profiles of Mtb strains can be available upon request. Whole genome sequence data generated in this work can be found under the BioProject accession number PRJNA1363199.

## Acknowledgments

We thank members of the Baliga lab, including Jake Valenzuela, Min Pan, Amardeep Kaur, and Julie Do for their critical discussions, feedback, and general lab resource support. We also thank the Aitchison lab for additional discussions and feedback. We thank Sean M. Gibbons for all of his feedback and guidance on the interpretation of the results in this manuscript. We thank Catherine Vilchèze and William Jacobs Jr. at the Albert Einstein College of Medicine for providing their *M. smegmatis* mc^2^155 wildtype, Δ*mshA* knockout, and pMV361::*mshA* complemented strains used in this study. We thank the Bhatt lab from Stanford University for providing us with the parental *M. smegmatis* mc^2^155 strain used in this study and from which evolved strains in this study were derived from. We thank the Molecular and Cell Core Facility of the Institute for Systems Biology. We acknowledge the authors of (S. Li et al., 2022) for their permission to use their data in our analysis. The article is licensed under a Creative Commons Attribution 4.0 International License (http://creativecommons.org/licenses/by/4.0/) and no changes were made to the data used. Lastly, we thank the Molecular Engineering and Sciences Institute at the University of Washington for their support.

This work was supported by National Institutes of Health grants R01AI141953 and R01AI128215, with additional funding from the Bill and Melinda Gates Foundation INV-009322 and INV-056403. This work was also supported with funding from the Tuberculosis Research Unit (TBRU) at the University of Washington (U19AI162583).

## Author Contributions

E.P.-T. contributed conceptualization, data curation, formal analysis, investigation, methodology, software, validation, visualization, and writing – original draft preparation (primary). V.S. contributed conceptualization, investigation, and methodology. F.D.M. contributed conceptualization, methodology, resources, and software. S.L. contributed investigation. S.R. contributed investigation and software. W.H. contributed investigation. A.D.Z. contributed conceptualization, investigation, and methodology. W.W. contributed software. M.S. contributed data curation and resources. D.T.M.H contributed data curation. S.J.D. contributed resources and supervision. T.N.T.T. contributed funding acquisition and data curation. S.T. contributed investigation, resources, and software. J.D.A. contributed funding acquisition and supervision. M.L.A.-O. contributed conceptualization, funding acquisition, methodology, resources, and supervision. N.S.B. contributed conceptualization, funding acquisition, methodology, project administration, resources, supervision, visualization, and writing – original draft preparation (support). Lastly, all authors contributed to writing – review and editing.

## Notes

https://github.com/evanpepper/Mycobacterium_PreR

### Competing Interest Statement

The authors have declared no competing interest.

