## Supplemental Figures for "Host oxidative stress primes mycobacteria for rapid antibiotic resistance evolution"

Pepper-Tunick et. al.

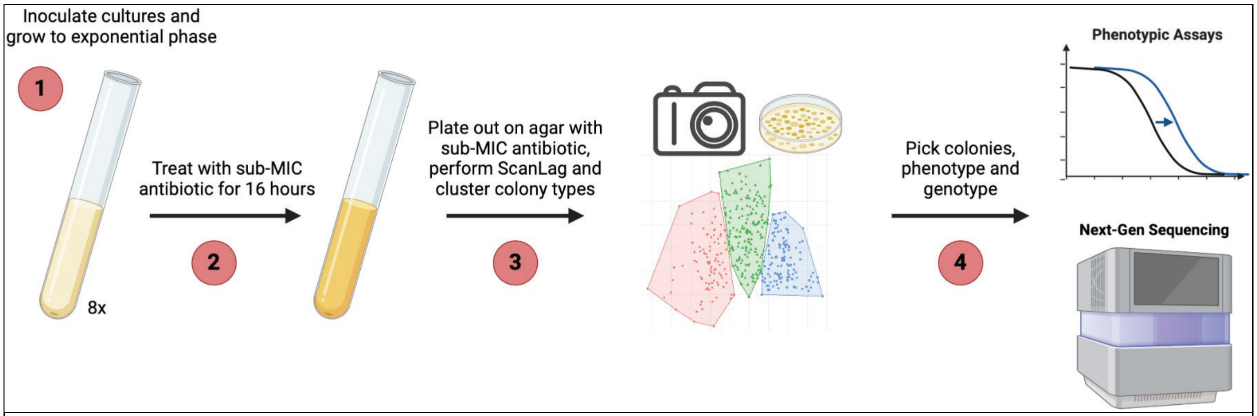


**Figure S1: A subinhibitory antibiotic treatment to enrich for pre-resistant mutants. (1)** Biological replicates of Msm mc$^2$155 cultures reach log phase ($OD_{600}$ 0.6–1.0). **(2)** Replicate cultures are $OD_{600}$ normalized into fresh liquid media and 2× $IC_{50}$ INH is added. Cultures are incubated at 37ºC for 16 hours **(3)** Cultures are plated out on solid 7H10 agar either with or without 2× $IC_{50}$ INH and ScanLag is performed. ScanLag data is then processed and colonies are clustered based on their growth characteristics. **(4)** Representative colonies from each cluster are then picked, grown isogenically, phenotyped for shifts in their $IC_{50}$, and sequenced using whole genome sequencing technologies to identify mutation(s) that confer the corresponding phenotype.

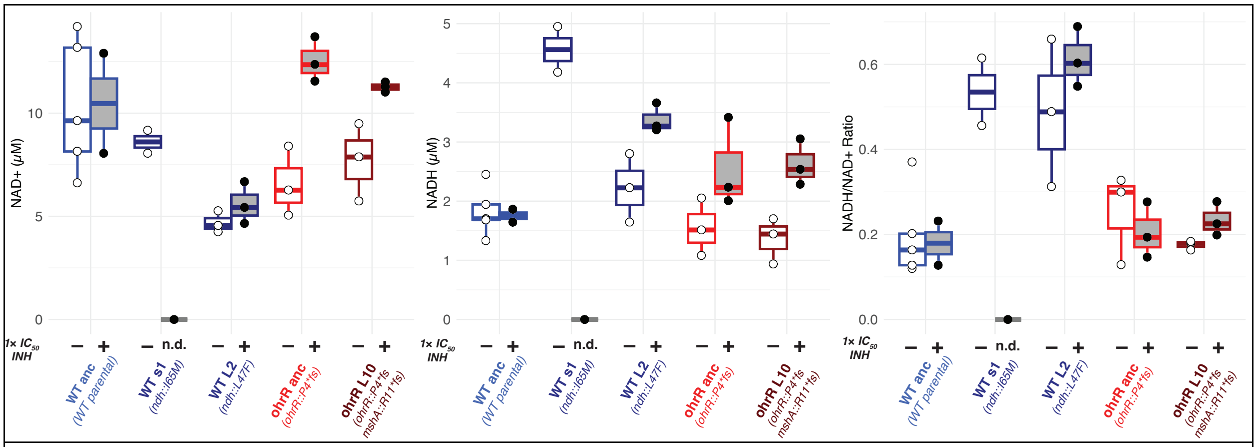


**Figure S2:** NAD+ quantification, NADH quantification, and NADH/NAD+ ratio during log phase growth of the wildtype, wildtype-derived INH$^R$ strains s1 (*ndh::I65M*) and L2 (*ndh::L47F*), *ohrR::P4*fs* LLRT strain, and its derived INH$^R$ strain L10 (*ohrR::P4*fs – mshA::R11*fs*) in the absence (indicated with "(–)") and presence (indicated with "(+)") of 1× $IC_{50}$ INH.

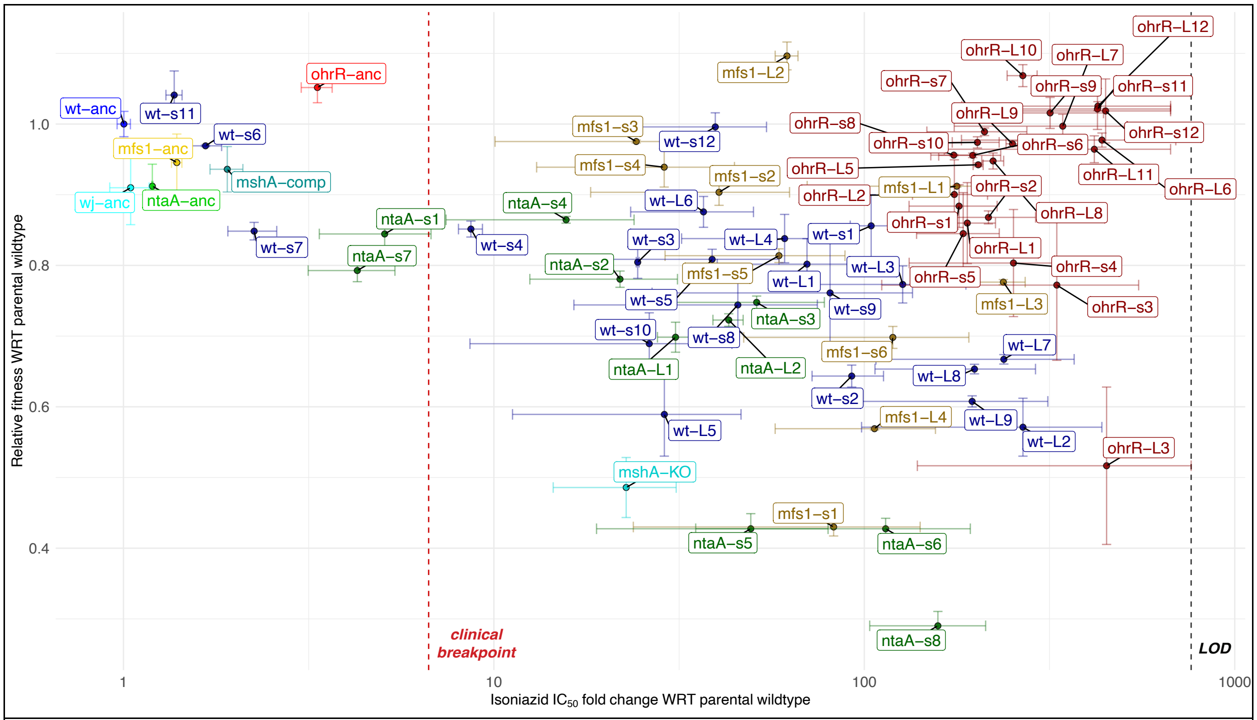


**Figure S3:** Various strains shown were grown to log phase and phenotyped in a dose response assay to quantify the INH $IC_{50}$ fold change and relative fitness with respect to the wildtype strain which they were originally derived from. Strain labels: wt-anc: Nitin Baliga Lab wildtype Msm mc$^2$155; wj-anc: William Jacobs Lab wildtype Msm mc$^2$155; *mshA*-KO: Δ*mshA* knockout derived from W.J. wildtype; *mshA*-comp: Δ*mshA* pMV361::*mshA* complemented strain derived from W.J. wildtype; mfs1-anc: *mfs1::G105D*; ntaA-anc: *ntaA_5::E50*fs*; ohrR-anc: *ohrR::P4*fs*. All remaining strains were derived from fluctuation assay of wildtype or LLRT strains.

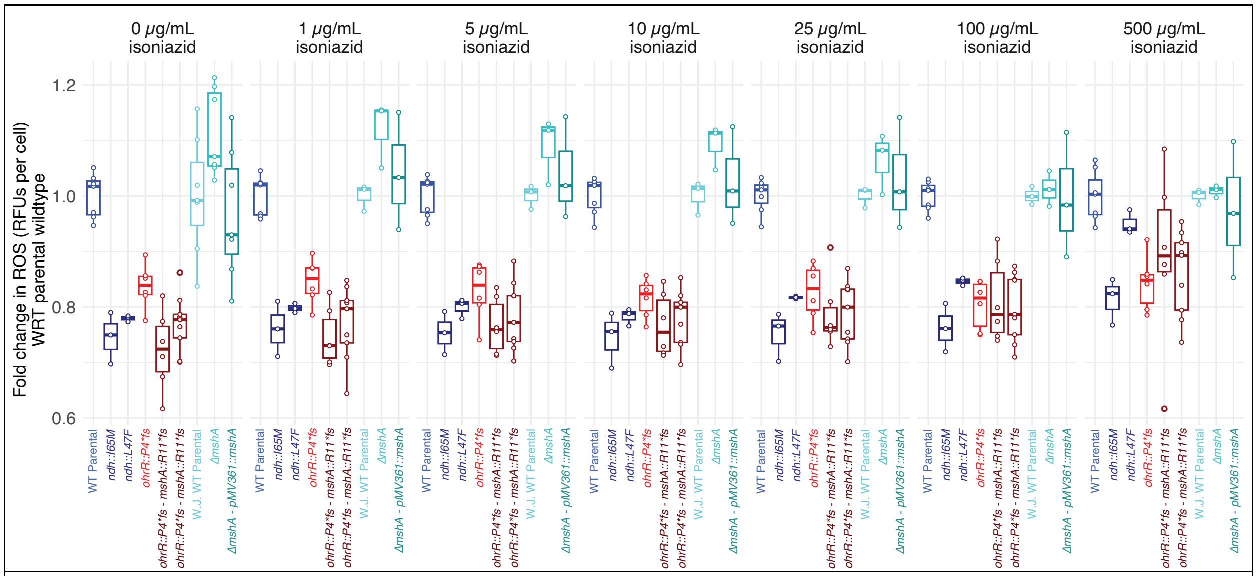


**Figure S4:** Various strains shown were grown to log phase ($OD_{600}$ 0.6–1.0) and then normalized to 8.0 x 10$^6$ cells / mL before exposing them to the fluorescent dye 2',7'-dichlorodihydrofluorescein diacetate (H2DCFDA), with or without INH at the specified concentration. RFUs were measured after 1 hour of incubation at 37ºC and normalized back to the cell count. Fold change RFUs for each strain is the ratio of the RFUs for a given sample relative to the RFUs of the wildtype strain from which each strain was derived, under the same conditions.

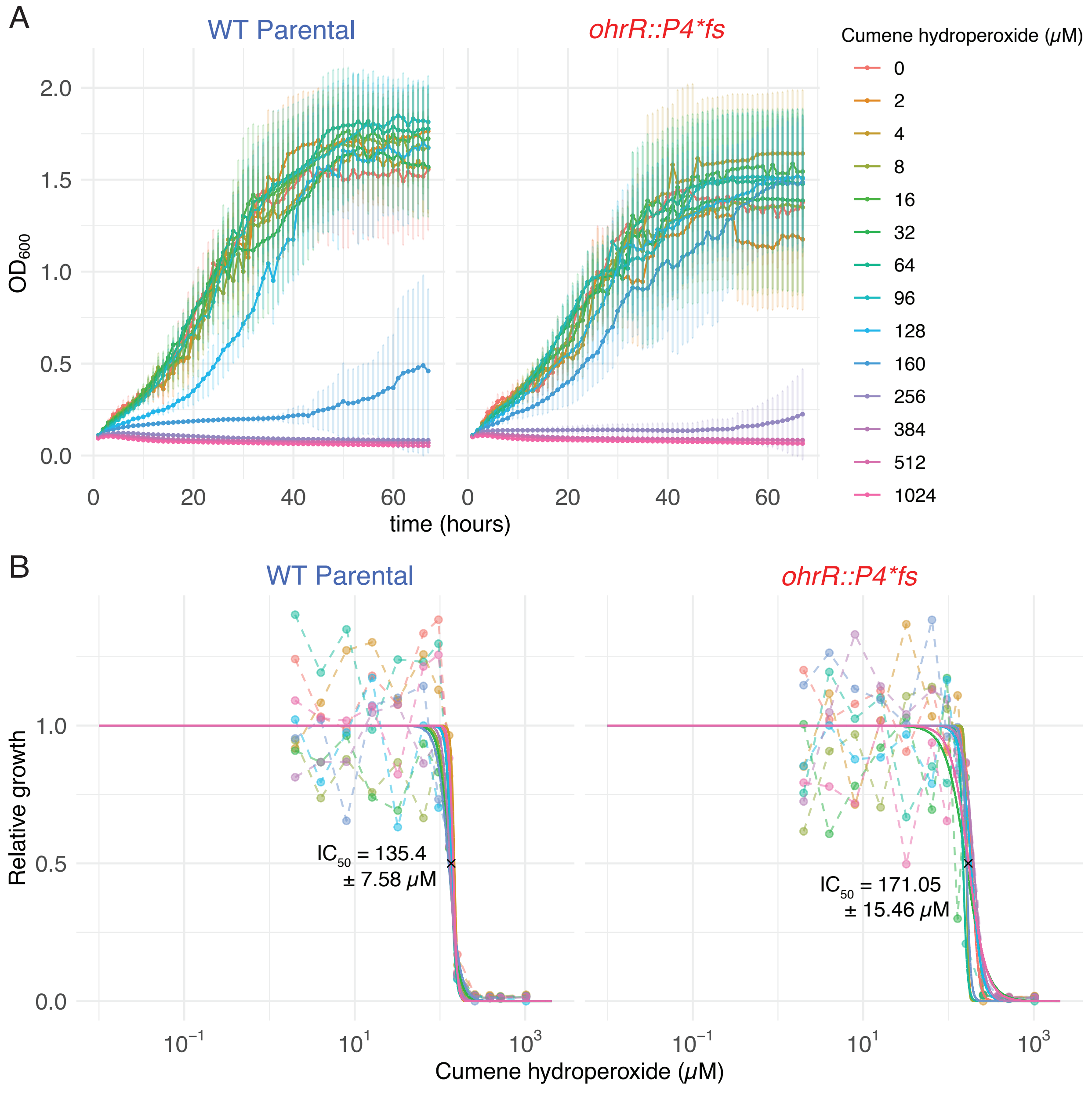


**Figure S5:** Dose response assay of wildtype and *ohrR::P4*fs* in cumene hydroperoxide. Strains shown were grown to log phase ($OD_{600}$ 0.6–1.0) and then normalized to $OD_{600}$ 0.02 before subjecting to a dose escalation of cumene hydroperoxide. $OD_{600}$ was measured over 48 hours of growth. Growth curves and dose response curves were generated as described in the Methods.
